# Competition between induced fit- and conformational selection binding: how different flux-based approaches help clarifying their interplay

**DOI:** 10.1101/2023.06.29.547035

**Authors:** Georges Vauquelin, Dominique Maes

**Affiliations:** Department Molecular and Biochemical Pharmacology, Vrije Universiteit Brussel, Pleinlaan 2, B-1050 Brussels, Belgium; Structural Biology Brussels, Vrije Universiteit Brussel, Pleinlaan 2, B-1050 Brussels, Belgium

**Keywords:** Induced fit, Conformational selection, Thermodynamic cycle Kinetics, Fluxes Simulations

## Abstract

**Background:** Binding kinetics has become a popular discipline in pharmacology. Yet, it was recommended a decade ago to look on ligand-target binding in terms of fluxes instead of rate constants. Like product formation in enzymology, these binding fluxes stand for the rate/velocity by which the target converts form one state into another. Several thereon-based approaches can be used for studying the functioning binding mechanisms.

**Objectives:** Although the relative contribution/importance of the induced fit and conformational selection binding pathways within a thermodynamic cycle is still hotly debated, only a single flux-based approach is customarily addressed for such calculations. There is a need to better understand the relevance of this peculiar approach.

**Methods:** The present findings relied on differential equation-based simulations, using the provided forward- and reverse rate constants for four reported cases as input.

**Results:** Both pathways may act in unison and even in tandem rather than being mutually exclusive. Also, calculating their relative contribution by distinct flux-based approaches may yield different outcomes for some of the cases under pre-equilibrium as well as equilibrium conditions. Additional flux-based approaches offer a rationale for the occurrence of those disparities.

**Conclusions:** The interplay between the induced fit- and conformational selection pathways is even more intricate than hitherto expected. It may me preferable to quantitate their relative contribution in terms of target occupancies instead of binding fluxes when genuine equilibrium can be reached.

**Perspectives:** Compared to the often-used elaborate algebraic equations, combining different binding flux-based approaches may offer better intuitive and visual insight into the functioning of complex binding mechanisms. More attention should also be paid to physiologically more relevant non-equilibrium situations.

## 1. Introduction

The duration and efficacy of a drug’s clinical action is governed by its pharmacokinetic (i.e. what the body does to drug) and pharmacodynamic (i.e. what the drug does to the body) properties. Those disciplines have long been considered to be complementary when dealing with reversible drug-target binding. While pharmacokinetics sufficed to explain the duration of its clinical effect, pharmacodynamic screening studies aimed to optimize drug candidates in terms of their efficacy and potency/affinity only. In this respect, experimental observations have meanwhile shed light on the non-negligible impact of the residence time of a ligand/drug (L) at its target (T) on this duration. Yet, the pioneering articles by Swinney et al. and Copeland et al. [1,2] and many ensuing reviews highlighted the clinical benefit of a long residence time (i.e. slow drug-target dissociation) and, in its wake, also on the thereto-beneficial contribution of induced fit (IF) binding. While this mechanism is likely to embody multiple steps [3–6], it is commonly referred to in terms of only two steps for the sake of simplicity; i.e. a fast reversible initial binding to yield a transient TL complex and a subsequent slow conformational transition thereof into a more stable T*L species [7–11]. Yet, T*L can also be reached via the conformational selection (CS) pathway according to which a conformational transition of T to T* precedes a fast and highly selective ligand binding step [10–15].

CS and IF binding mechanisms have often been regarded to be mutually exclusive [11,16–18]. In line with this viewpoint, Copeland [19,20] concluded about 10 years ago that most of the drugs with high clinical efficacy act via IF. Yet, this explicit distinction between IF- and CS-binders is now increasingly abandoned in favor of a 4-state thermodynamic cycle model in where both mechanisms act alongside as pathways for progressing till T*L (Figure 1) [16,21–25]. This updated viewpoint largely benefited from the introduction of a novel binding flux, (F), -based approach by Hammes et al. [22]. Alike the rate of product formation in enzymology, a binding flux refers to the rate by which a given target species converts into another one (e.g. from T to TL, Figure 1). They argued that the relative contribution of the IF and CS pathways to T*L should be determined by comparing their “macroscopic” forward fluxes that account for the overall conversion of T into T*L (F_on,_ Table 1) rather than by merely comparing rate constants. Based on this approach, they observed that the relative contribution/importance of IF (Rc_-IF_, Table 1) increases with the ligand concentration, [L] at equilibrium.

**Figure 1.**
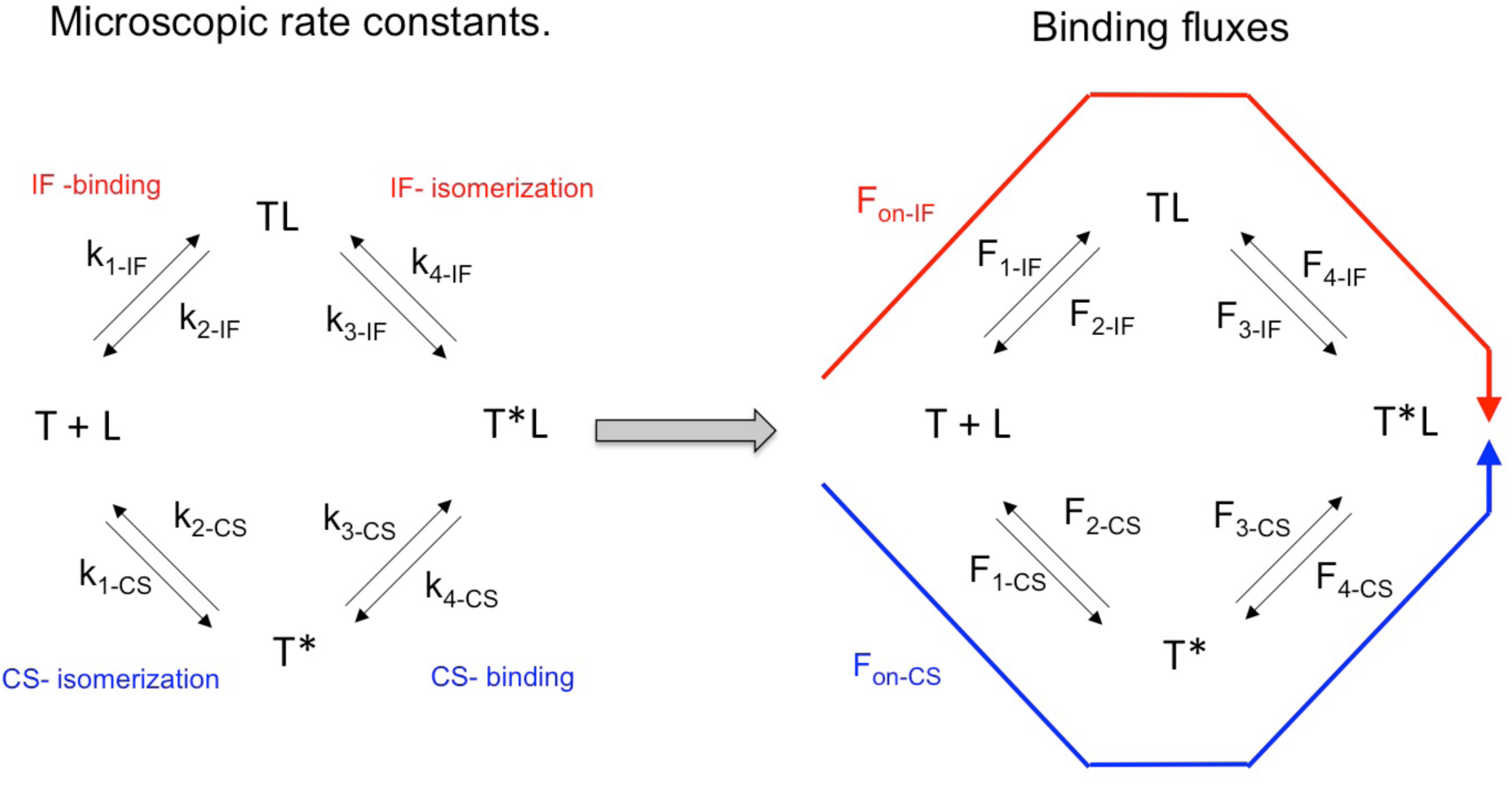
Nomenclature for an IF – CS-based thermodynamic cycle. L Is the ligand, T and TL are targets in the “native” conformation; T* and T*L are targets in the ”active” conformation. The nomenclature for the microscopic fluxes is in accordance with the nomenclature for the related rate constants [39]. The macroscopic forward, and reverse fluxes (i.e. F_on_ and F_off_) for each pathway refer to the overall rate by which T converts into T*L and vice versa.

**Table 1.**
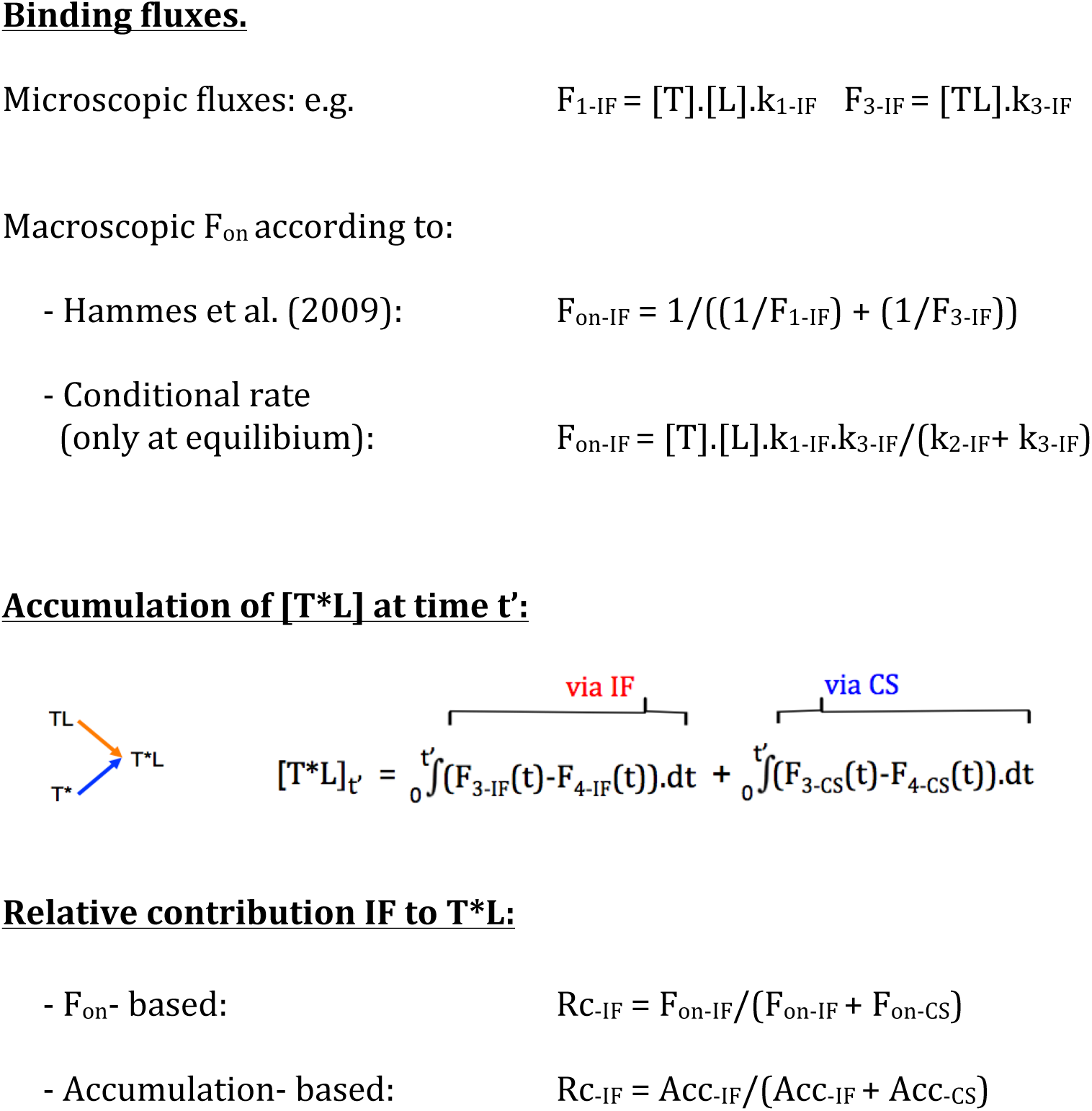

Meanwhile, several others have consistently reproduced this pattern [16,17,26–31]. Equations have also been elaborated to express this relationship [17,24,31] and the current picture is that IF becomes fully dominant at high [L] whereas CS usually dominates at low [L], but not always [27]. The observed variance of Rc_-IF_ at low [L] has been related to merely two rate constants [31]. Interestingly, Ordabayev et al. [29] recently presented an alternative and, a potentially even more attractive approach to quantitate Rc_-IF_, i.e. by comparing by how much both pathways did contribute to the accumulation of [T*L] till equilibrium (Table 1). Although both approaches are quite distinct from the conceptual point of view, they provided relatively comparable Rc_-IF_ - [L] relationships for the hitherto documented examples [29,3,31]. However, it is of note that those examples largely complied with the classical definition of CS, which stipulates that T* represents a thermodynamically unstable high-energy conformation so that it only constitutes a minute fraction of the total target population in the absence of ligand. Yet, this is not always the case for some targets. Indeed, the presence of co-existing unbound target states was already detected by X-ray crystallography. Recent advances in molecular dynamics simulations and in experimental techniques like fluorescence- and nuclear magnetic resonance spectroscopy revealed that a sizable fraction of the unbound target population may also adopt T*-like conformations under more relevant physiological conditions [28,32–37]. Moreover, those advances also led to the detection of population shifts between T* and T*L in the presence of distinct ligands. It is of note that such shifts are usually interpreted in terms of CS [38].

Those assertions prompted us to gain a better insight into how the presence of an initial sizable [T*] may affect ligand binding to an IF - CS-based thermodynamic cycle. Relying on an extended set of still little exploited binding flux-based approaches; we here focused on how the individual steps and pathways progress to the overall, macroscopic equilibrium. The thereto-dedicated simulations are based on the rate constants that were reported in the literature for four distinct cases. Present findings do shed light on the sometimes-intricate interplay between both pathways and on the fact that the distinct approaches for calculating the contribution of IF and CS to the ultimate T*L complex may produce different outcomes. This prompted us to critically reflect on the most adequate approach for quantitating the Rc_-IF_ for such thermodynamic cycle.

## 2. Materials and Methods

### 2.1. Nomenclature and definitions

Square brackets refer to concentrations. The “microscopic” fluxes for individual steps are denoted as F_1_, F_2_… in accordance with the nomenclature for the related rate constants [39] (Figure 1). CS- and IF-appends refer to the pathway in question. Such microscopic fluxes correspond to the product of the concentration of the starting target species and either with the associated first-order rate constant (i.e. for a mono-molecular step) or with the associated second-order rate constant and [L] (i.e. for a bimolecular ligand binding step). The published “microscopic” rate constants for each of the investigated cases are provided and commented upon in Supplementary Table S1.

The macroscopic “forward” (F_on_) and the “reverse” fluxes (F_off_), for each pathway refer to the overall conversion of T into T*L and backwards. Rc_-IF_ stands for the relative contribution of the IF pathway to T*L within a thermodynamic cycle. This parameter is traditionally based on F_on-IF_ and F_on-CS_ such as shown in Table 1 [22]. IF becomes dominant when Rc_-IF_ exceeds 0.5 and the relative contribution of the CS pathway to T*L equals 1-Rc_-IF_. Rc_-IF_ can also be calculated based on other criteria such as by how much the IF-isomerization-(earlier denoted by us as transconformation) and the CS-binding steps contribute to the accumulation of [T*L] (Table 1) [29–31].

### 2.2. Simulations and calculations

F_on_- and F_off_-values of each pathway can be related to their constituent microscopic fluxes at any time point. This implies prior knowledge of [L], the eight microscopic rate constants of the cycle. The concentration of the four target species at any time point can be obtained by integrating in parallel the differential equations given in Supplementary Table S2. In practice, such solutions were provided by consecutively solving the equations over very small time intervals; i.e. by Euler’s method [40,41]. Such concentrations can also be obtained by making use of e.g. the KinTek Explorer^®^ software (Kintek, Snowshoe PA).

Target concentrations at the onset take account of a pre-existing equilibrium between [T] and [T*]. Supplementary Figure S3 provides a step-by-step account of how [T] and [TL] can be converted to the macroscopic F_on_ of the IF pathway. F_on_- and F_off_ values can also be conveniently calculated by using conditional rate-based equations that only require [T], the microscopic forward rate constants and [L] as input (Table 1) [24,42]. Yet, this method only applies to equilibrium binding [31].

Differential equations may also reveal by how much the concentration of each target species has been modified by each of the therewith-connected microscopic steps till a given time point. In this respect, it is of note that those microscopic steps permit one target species to accumulate at the expense of an equivalent de-accumulation of the therewith-connected species. The arrows that designate those steps in Figures 5C and D and 6B to D point in the direction of the target species that (at least initially) accumulates. The therewith-corresponding differential equations are given in Supplementary Table S2. Although the present arrows point mostly rightward, they can occasionally also point leftward such as illustrated for the CS-isomerization step in Figure 6B to D. The contribution of the IF-isomerization- and of the CS-binding step to the time-wise accumulation of [T*L] can be obtained by integrating equations 7 and 8 in Supplementary Table S2.

**Figure 5.**
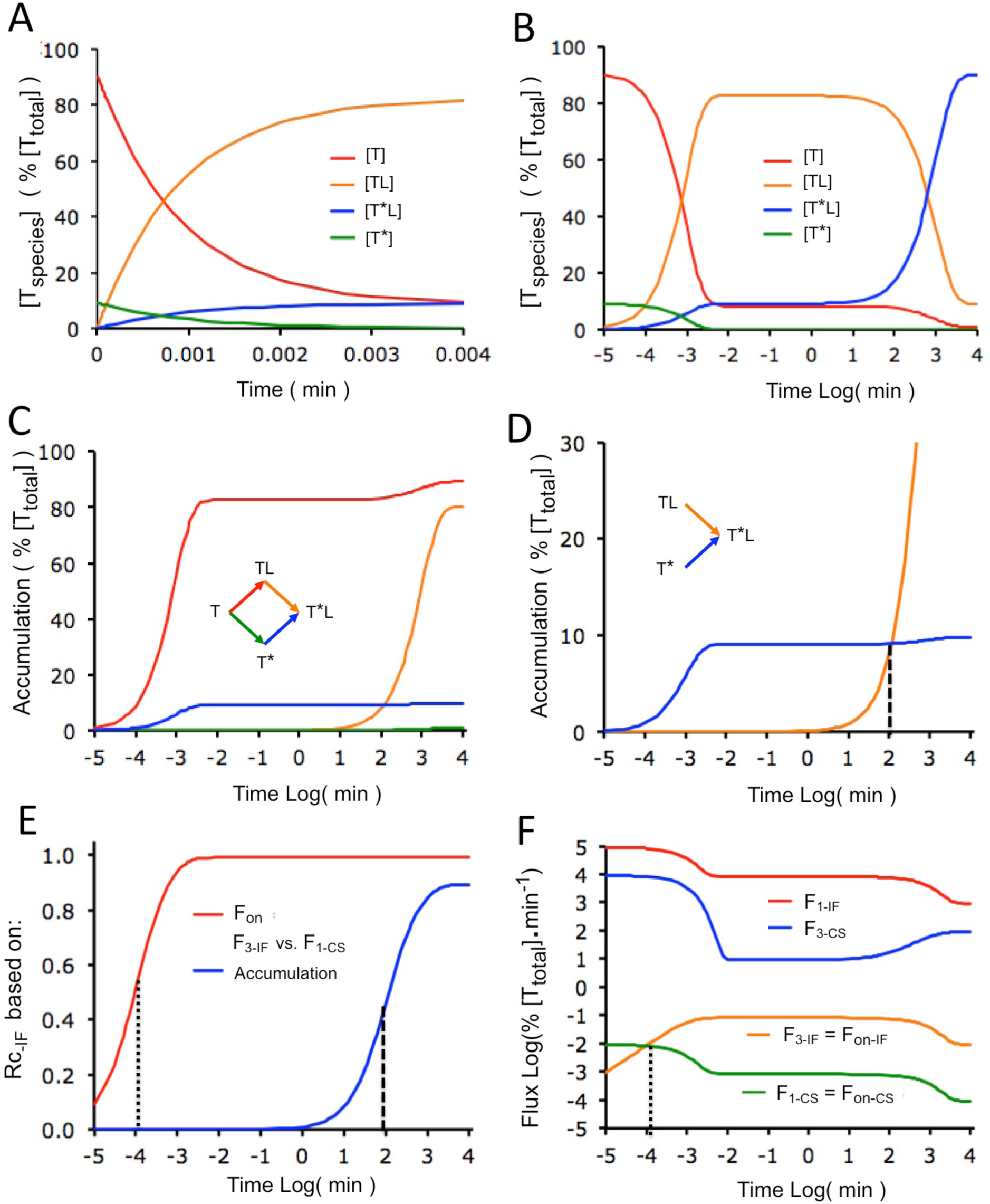
Closer scrutiny of Case C. A and B) Time-wise evolution of the concentration of each target species after adding 10 µM of ligand (= 100 x the K_D_ of the cycle). Panel A focuses on the fast initial binding of the ligand to T and T*. The logarithmic time scale of panel B provides an overview of all evolutions. Please note the slow conversion of almost all TL into T*L. C and D) Time-wise accumulation of each target species via the connected microscopic steps (arrows, please see section 2.2 for more details). Panel C provides an overview for all target species. Please note that the sizable [TL] at equilibrium in Paned B amounts to the difference between the final [TL] (red curve) and [T*L] (orange curve). Panel D focuses on the moment when the accumulation-based Rc_-IF_ equals 0.5 (large dots): i.e. when the contribution of the IF-isomerization step equals the earlier contribution of the CS-binding step. E) Evolution of F_on_- and accumulation-based Rc_-IF_ values with time. Dots indicate the moment when each Rc_-IF_ value equals 0.5, such as outlined for Panels D and F. F) Double-log plot for the time-wise evolution of the four microscopic forward fluxes as well as of F_on-IF_ and F_on-CS_. Those macroscopic fluxes closely equal the smallest of their constituent microscopic fluxes; i.e. F_1-IF_ and F_3-CS_, respectively. The F_on_-based Rc_-IF_ equals 0.5 at the moment when F_1-IF_ and F_3-CS_ cross (small dots).

**Figure 6.**
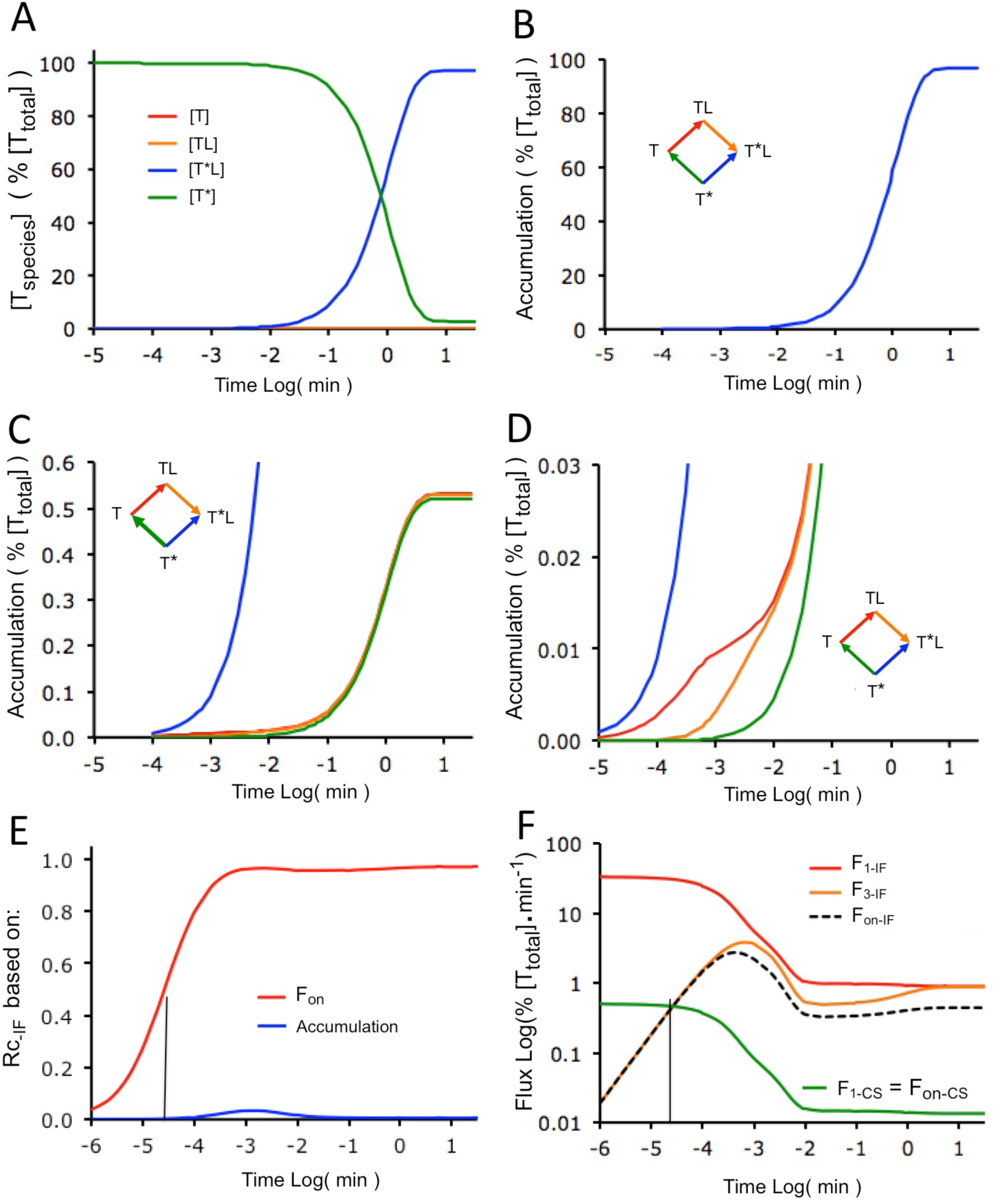
Closer scrutiny of Case D. A) Time-wise evolution of the concentration of each species after adding 30 μM of ligand (= 3.3 10^5^ x the K_D_ of the cycle). B to D) Time-wise accumulation of each target species via the connected microscopic step (arrows). The outspoken contribution of the CS-binding step to the to accumulation of [T*L] (blue arrow) is clearly evidenced at the largest scale (B). The medium scale (C) unveils additional, nearly overlapping CS-reverse isomerization-(inversed green arrow), IF-binding and IF-isomerization mediated accumulations of T, TL and T*L, respectively. This plot indicates that about 0.5 % of the final [T*L] originates from T* via a fast succession of those steps. The smallest scale (D) unveils an additional, even faster succession of IF-binding and IF-isomerization mediated accumulations of TL and T*L. This plot indicates that the initial [T] (0.01 % of [T_tot]_) is converted into T*L via IF. E) Evolution of F_on_- and accumulation-based Rc_-IF_ values with time. The accumulation-based Rc_-IF_ remains minimal throughout. The solid line indicates the moment when the F_on_-based Rc_-IF_ equals 0.5. F) Double-log plot for the time-wise evolution of the three smallest microscopic fluxes, F_on-IF_ and F_on-CS_. The F_on_-based Rc_-IF_ equals 0.5 when F_1-IF_ and F_3-CS_ cross (solid line).

### 2.3 paradigms

#### 2.3.1. Essential paradigms

- The simulations are based on the conventional paradigm that the free ligand concentration, [L], largely exceeds the total target concentration, [T_tot_], so that it remains constant with time.

- The IF- and CS- pathways have to produce the same difference in Gibbs free energy between T and T*L. This “detailed balance rule” dictates that the thermodynamic equilibrium dissociation constant, K_D_, values of those pathways (i.e. their k_2_.k_4_/k_1_.k_3_ ratios) must be equal [24].

#### 2.3.2. Paradigms for a classical cycle

- [T*] represents a sparsely populated high-energy conformation. Accordingly, this target species only constitutes a minor fraction of [T_tot_] at equilibrium in the absence of ligand. In kinetic terms, this requires k_2-CS_ to largely exceed k_1-CS_.

- The “rapid equilibrium” paradigm requires the ligand-binding step to proceed faster than the isomerization step for each pathway [26,43]. In kinetic terms, this requires k_2-IF_ to exceed k_3-IF_ for IF [44] and k_4-CS_ to exceed k_1-CS_ for CS [23,43].

## 3. Results

### 3.1. Ligand saturation binding and association plots

Figure 2 compares the saturation binding plots of four different cases that have been reported in the literature. The microscopic rate constants that are necessary for the present simulations are provided in Supplementary Table S1. Those constants result from nuclear magnetic resonance measurements of the binding of NADPH to dihydrofolate reductase [21,22] for case A, molecular dynamics simulations of MDM2 binding to the intrinsically disordered transactivation of the tumor suppressor p53 protein [27] for case B, a theoretical example [26] for Case C and the binding of flavin mononucleotide cofactor to *Desulfovibrio desulfuricans* flavodoxin [22] for case D. Of note is that Case A and D constitute the two examples that were documented by Hammes et al. [22]. The plots in Figure 2 depict the concentration of the two ligand- bound target species (i.e. [TL] and [T*L]) for each cycle at equilibrium for different [L]. Two observations merit attention. First, while [T*L] largely exceeds [TL] for cases A and D, the latter already constitutes a sizable portion of the binding for case C and it even exceeds [T*L] for case B at all [L]. Those differences stem fact that the [TL]/[TL] ratio corresponds to the k_4-IF_/k_3-IF_ ratio (provided in Supplementary Table S2) at equilibrium. Second, while there is reasonable concordance between [L] at which [T*L] is half-maximal and the thermodynamic K_D_ of the cycle (i.e. the k_2_.k_4_/k_1_.k_3_ ratio) for the three first cases, case D is clearly an outlier.

**Figure 2.**
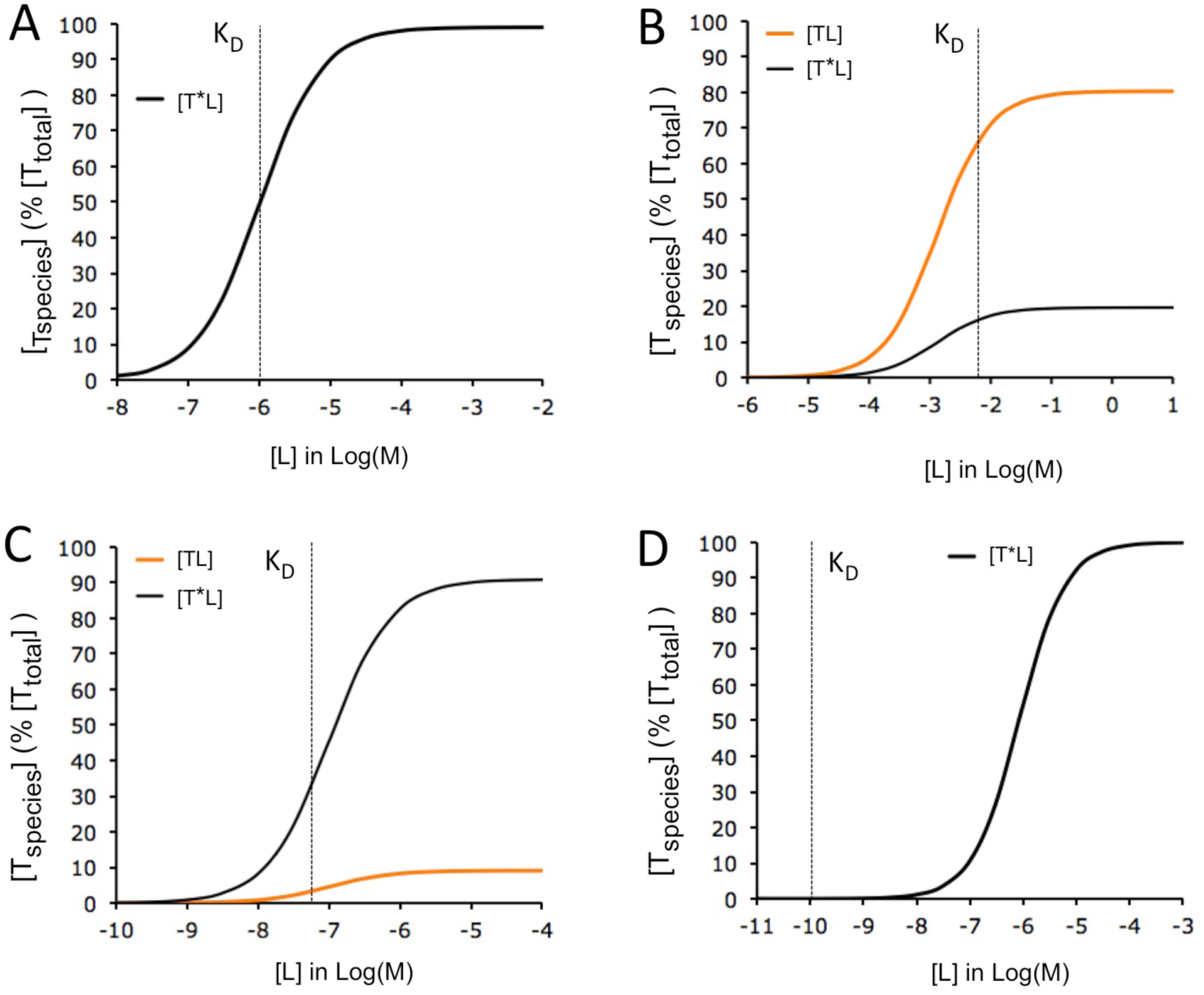
Saturation binding curves for cases A to D. Presented Cases: A) Example 1 in [22] B) from [27] C) from [26], D) Example 2 in [22]. The plots show how the equilibrium- concentration of each target species of interest varies with [L]. Only the species whose concentration is non-negligible are shown. The plots are based on the microscopic rate constants that were provided in the literature (listed an commented upon in Supplementary Table 1). K_D_ is the thermodynamic equilibrium dissociation constant (i.e. k_2_.k_4_/k_1_.k_3_) for each pathway. This parameter must be equal for both pathways of the same cycle.

Figure 3 compares the time wise variations of [TL], [T*L] and [T*] for the cases above. [L] Is deliberately kept high to comply with the fact that optimal drug therapy often requires high drug concentrations near their targets [45]. Moreover, such simulations comply more with the conventional “rapid equilibrium”- and “unchanging free ligand concentration” paradigms. IF binding has repeatedly been reported to exhibit a transient [TL]-“overshoot” under such conditions (e.g. [30,46,47]). Such overshoots are also observed for cases A to C in Figure 3 and the positive relationship between the [TL]- maximum and [L] is further illustrated for case A in Supplementary Figure S4. In all instances, [TL] rises very swiftly and declines only slowly afterwards. This decline is usually related to a slow conversion of the [TL]- excess of into [T*L]. Yet, this interpretation does not always apply for a IF- and CS- based thermodynamic cycle since it was recently demonstrated that such overshoot can also be resorbed via the CS-pathway when the T ➞ T* transition is sufficiently fast [47]. Case D represents a peculiar situation in where the equilibrium between [T] and [T*] permits the latter to nearly fully account for the free targets at the onset of the binding simulations. Rather than the expected [TL]- overshoot, we here merely observe a slow drop of [T*] along with an equally slow rise of [T*L], thereby already suggesting that the vast majority of the binding proceeds via CS.

**Figure 3.**
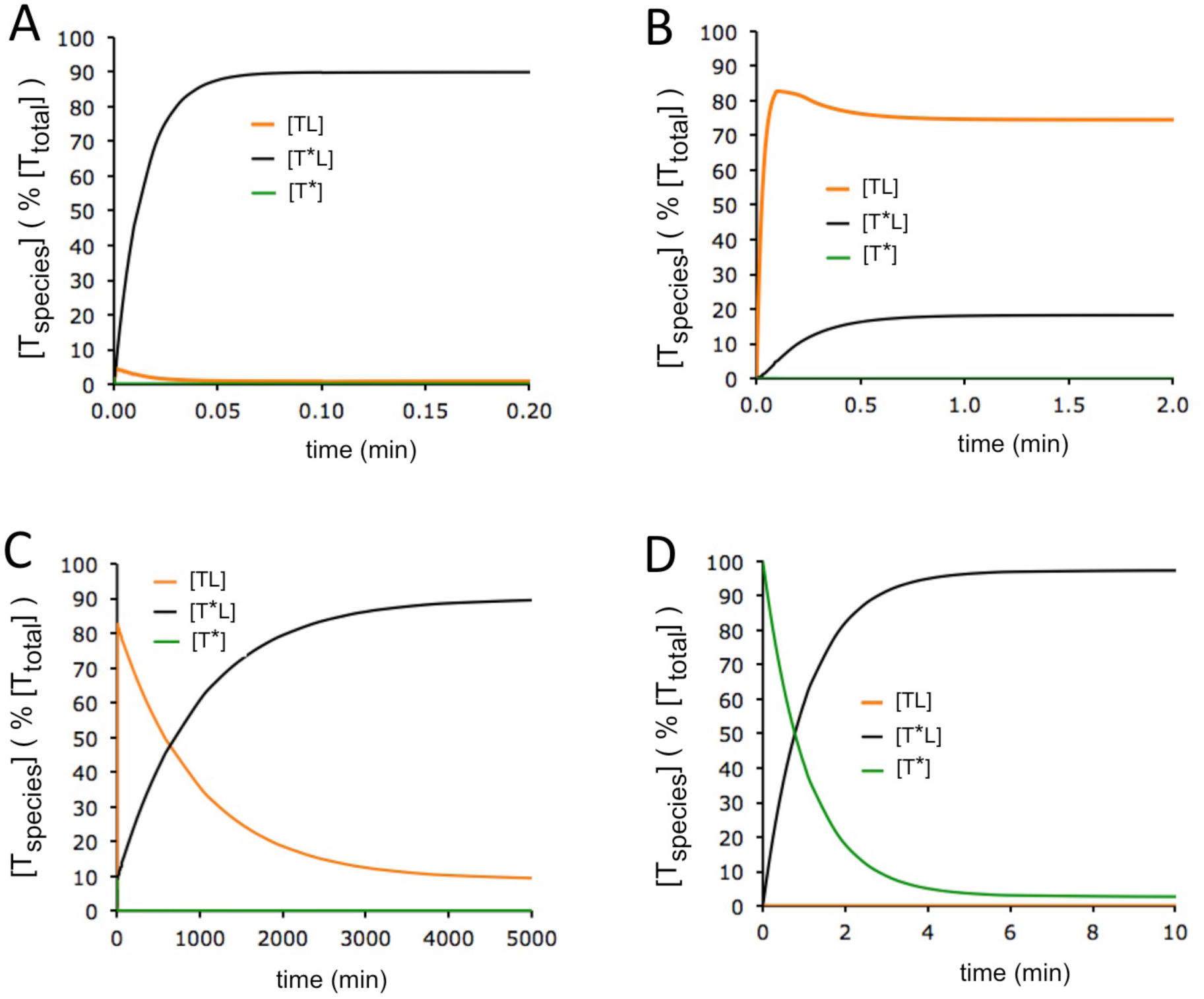
Ligand association curves for cases A to D. The plots show how the concentration of each target species of interest varies with time at high [L]. A) [L] Equals 10 µM (= 10 x K_D_) for case A, 16.5 mM (= 2.5 x K_D_) for case B, 10 µM (= 100 x K_D_) for case C, 30 mM (= 3.26 x 105 x K_D_) for case D.

Supplementary Figure S5 further compares the time-wise evolution of the observable binding of L in e.g. radioligand association experiments (i.e. [TL] + [T*L]) with potential target “activity” (i.e. [T*] + [T*L]) in functional experiments for the four cases. However, this latter conformation has also been evoked in the context of antagonist binding [48,49], so that, in general, it should merely be considered to be distinct from that of the ground state, T,

### 3.2. F_on_- and accumulation- based Rc_-IF_ profiles

Figure 4 compares F_on_- and accumulation- based Rc_-IF_ vs. [L] plots for these four cases. An explicit account of mode of calculating such Rc_-IF_ values is provided in section 2.2 and Supplementary Figure S3. Those data refer to equilibrium binding for the sake of comparison as well as to remain in tune with the prevailing practice. Several observations merit special attention. First, the F_on_- based Rc_-IF_ values vary considerably from one case to another at very low [L], i.e. from nearly baseline for case C till 0.99 for case B. In this respect, it is of note that F_on_-based Rc_-IF_ values can be related to some of the microscopic rate constants when [L] = 0 [31].

**Figure 4.**
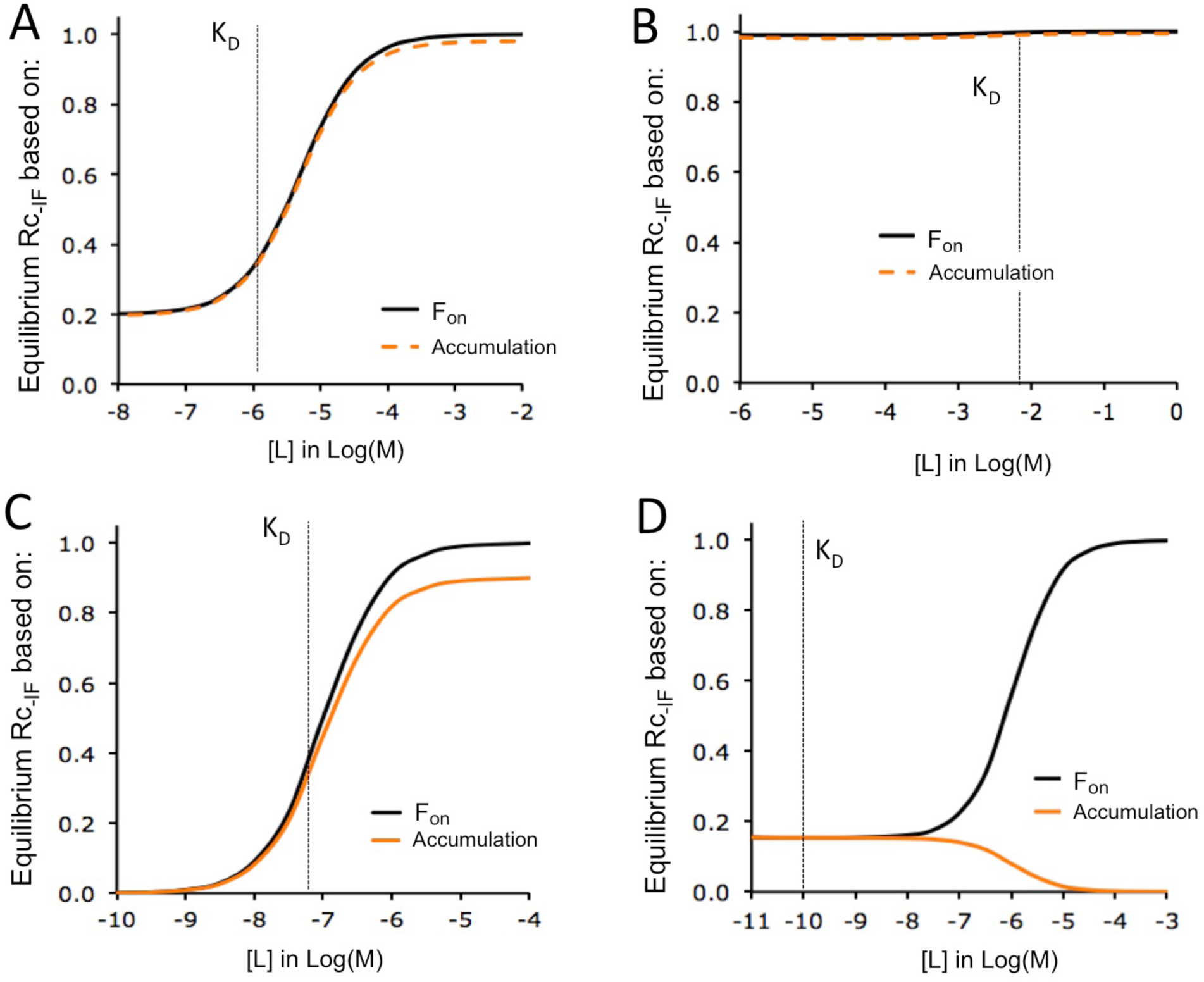
Equilibrium- Rc_-IF_ versus [L] plots based on the F_on_- and [T*L]- accumulation for cases A to D. Rc_-IF_ stands for the relative contribution of IF to [T*L]. The corresponding equations are provided in Table 1. The relative contribution of CS equals 1- Rc_-IF_. Both pathways contribute equally when Rc_-IF_ = 0.5.

Second, these four cases have in common that their F_on_-based Rc_-IF_ values increase till unity with [L]. In this respect, please note that the plots in Figure 4 in are very similar to those that were initially presented by the respective authors. Admittedly, this increase is only barely perceptible for case B due to the initially already exceedingly high Rc_-IF_ value. Also, the plot for case D clearly illustrates that such Rc_-IF_ values do not necessarily increase half- maximally when [L] equals the thermodynamic K_D_ of the cycle.

Third, the F_on_- and accumulation- based Rc_-IF_ versus [L] plots nearly overlap for cases A and B only (Panels 4A and B) and similar patterns have also been shown for earlier theoretical cases [30,31]. Such concordance is clearly unrelated to the magnitude of the minimal Rc_-IF_. However, it is of interest to note that all those simulations start from an equilibrium situation in the absence of ligand (i.e. in where [T*] is solely governed by the k_1-CS_/k_2-CS_ ratio), and that this initial [T*] is very low for all the cases that are mentioned above. Indeed, the initial [T*] only represents 1 % of [T_tot_] in previous work, 2 % for case A and merely 0.11 % for case B. It is thus possible that those overlaps are linked to a low initial [T*].

Examination of cases C and D (Panels 4C and D) lends support to this view. Both approaches give rise to dissimilar and even to diverging Rc_-IF_ - [L] plots for those cases and their initial [T*] is also distinctly higher. Indeed, [T*] already accounts for 10 % of [T_tot_] for case C and it is remarkable that its accumulation- based Rc_-IF_ vs. [L] plot is also about 10 % lower than its IF- based counterpart at all concentrations. Although it may be argued that case C is purely theoretical, concordant observations have been made for the activation of the UvrD helicase enzyme of *Esherichia coli* by its accessory MutL factor: i.e. a case whose initial [T*] was alike (shown in Supplementary Figure S6 for [29]). The F_on_- and accumulation- based plots may even diverge completely, such as shown by the descending profile of the latter for case D. Peculiar to this case is that (based on the k_1-CS_/k_2-CS_ ratio thereof) [T*] should amount no less than 99.99 % of [T_tot_] at the initial equilibrium.

Taken together, the present findings suggest that the initial [T*] is somehow related to the extent by which F_on_- and accumulation- based Rc_-IF_ vs. [L] plots are able to diverge. By making use of additional binding flux- based approaches we next scrutinize the mechanisms for that contribute thereto for cases C and D. Since the dissimilarity between those plots is most perceptible at high [L], we will further only focus on this condition.

### 3.3. Binding characteristics of case C at high [L]

Figure 3C already showed how the concentration of each target species of case C evolves with time for [L] = 10 µM. However, its extended linear time scale may preclude the detection and comparison of early important events. Figure 5A provides a more precise account of those. The plots suggest that the equally fast decrease of [T*] and equal increase of [T*L] owes to a fast CS- binding step. Similarly, a fast IF- binding step likely accounts for the equally fast decrease of [T] and equal increase of [TL]. The logarithmic time scale of Figure 5B (and ensuing Panels) is necessary for comparing the processes that take place at quite different timeframes. Besides those early events, Figure 5B also suggest that the equally slow decrease of [TL] and remainder increase of [T*L] owes to a slow IF- isomerization step.

The above suggestions are confirmed by comparing the contribution of the microscopic steps to the accumulation of each target species over time (indicated with arrows in Figure 5C). Indeed, the IF and CS- binding steps are very swift. The initial IF-binding step is followed by a much slower IF-isomerization step and (such as even better illustrated in Figure 5D) the CS-binding step only plays a modest further role later on. It is thus the very fast binding of the ligand to the already sizable initial [T*] (blue curve) that essentially prevents IF to exclusively contribute to the accumulation of T*L.

Figure 5E compares the time- wise progression of the F_on_- and accumulation- based Rc_-IF_ values till their final plateau. The slow progression of the latter can be related to the very slow IF-isomerization step. Indeed, Rc_-IF_ reaches the 0.5- mark when the IF and CS contribute equally to the accumulation of T*L. Such as shown in Figure 5D, this parity is presently attained when the contribution of the slow IF-isomerization step (orange curve) finally catches up with the prior contribution of the CS-binding step (blue curve).

By contrast, the much faster progression of the F_on_- based Rc_-IF_ in Figure 5E relies on flux-based criteria. In this respect, two basic principles need to be taken into account. First, the smallest of the microscopic forward fluxes within each pathway largely determines the F_on_ thereof. Figure 5F clearly shows that this role is conferred to the microscopic fluxes of the isomerization steps (i.e. F_3-IF_ and F_1-CS_). This equivalence is also evidenced by the close overlap between the F_on_- and F_3-IF_/(F_3-IF_ + F_1-CS_)- based Rc_-IF_ versus time plots in Figure 5E.

Second, the amplitude of a microscopic flux varies in par with the concentration of the target species it originates from (per definition). Accordingly, the early increase of F_on-IF_ is dictated by [TL] whereas the early decrease of F_on-CS_ is dictated by [T]. This implies that the F_on_- based Rc_-IF_ reaches the 0.5- mark early on (Figure 5E) because of the rapid binding of L to nearly all pre-existing T (Figure 5A and B) and the therefrom- resulting rapid decrease of [T] (and thus F_1-CS_) and increase of [TL] (and thus F_3-IF_). The F_on_- based Rc_-IF_ progresses also faster to its plateau than its accumulation- based counterpart for cases A and B for the same reasons (Supplementary figure S6).

Finally, it is important to verify that the simulated ligand association experiment lasts long enough to approximate equilibrium binding. This issue can be addressed by two distinct approaches. First, all the forward and reverse fluxes should have reached (or at least become very close to) unity. Such as shown by the forward/reverse flux ratios in Supplementary Figure S7, this is presently the case after a sufficiently long incubation. Second, genuine equilibrium- F_on_- based Rc_-IF_ values can be conveniently calculated by making use of conditional rate- based equations [24,31,42]. The terminating simulation- based values in Figure 5E also equal to the calculated ones.

### 3.4. Binding characteristics of case D at high [L]

Figure 6A depicts the time-wise evolution of each target species for [L] = 30 µM. The most striking observations are: the absence of sizable [T] throughout, the initially utterly high [T*] and a nearly full decline of [T*] along with a concomitant equivalent increase of [T*L]. Figure 6B reveals that this T-to-T*L conversion essentially owes to the CS-binding step. It is important to note that the F_on_-based Rc_-IF_ values are able to reach 1 at high [L] (Figure 4C). This observation suggests that IF must be operating as well. Focusing on manifestations of lesser amplitude shows that this is the case. Indeed, it first appears that the three other steps affect about 0.5% of [T_tot_] each (Figure 6C). More precisely, the reverse isomerization of T* contributes to the accumulation of T, ligand binding to T contributes to the accumulation of TL and the isomerization of TL contributes to the accumulation of T*L. In this respect please note that the arrow of the CS-isomerization step (in green) now points to the left. The fact that these three curves nearly overlap indicates that the concerned microscopic steps act in fast succession. When focusing on variations of even lesser amplitude (Panel 6D) it also appears that all initial T (i.e. about 0.01% of the total target population) is swiftly converted into TL and that the conversion of TL into T*L takes place at a lower pace. Albeit of minute amplitude, this pattern is typical for an IF- related [TL]- overshoot at high [L]. Taken together, the present observations reveal that most of the IF pathway is actually “fuelled” by conversion of a small fraction of the initial T* into T.

The accumulation- based Rc_-IF_ remains constantly very low in Figure 6E. This pattern reflects the paramount contribution of the CS-binding step to the accumulation of T*L such as shown in Figure 6B. By contrast, Figure 6E shows a rapid rise of the F_on_- based Rc_-IF_ till almost full IF- dominance. Such as for case C, closer inspection Figure 6F reveals that the early amplitudes of F_on-IF_ and F_on-CS_ are also largely dictated by F_3-IF_ (orange curve) and F_1-CS_ (green curve), respectively. F_3-IF_ rises also swiftly till well above F_1-CS_ early on and this situation persists till global equilibrium is reached. Taken together, the F_on_- based approach allows IF to already largely dominate early on in spite of the only minute contribution of this pathway to the accumulation of T*L for this peculiar case.

### 3.5. Impact of the initial [T*] on the equilibrium- Rc-IF values

The lower accumulation- based equilibrium Rc_-IF_ values for cases C and D is likely to be related to the presence of already sizable initial [T*]. The present simulations aimed to get gain better insight into this issue. Changing the initial [T] can easily be achieved by modifying k_1-CS_ and/or k_2-CS_. Yet, the “detailed balance rule” dictates that at least one additional rate constant should be changed as well. Since this would alter the kinetic properties of the cycle, we rather opted for exploring non-equilibrium situations at the onset: e.g. such as when the target molecules are formed in the basal T- conformation but that [T] – [T*]- equilibrium is not yet reached at the moment that the ligand is added.

Figure 7A and C show that increasing the initial [T*] (till its initial equilibrium- based value) hardly affects the F_on_- based Rc_-IF_. By contrast, it produces a proportional decrease of the accumulation- based Rc_-IF_.

**Figure 7.**
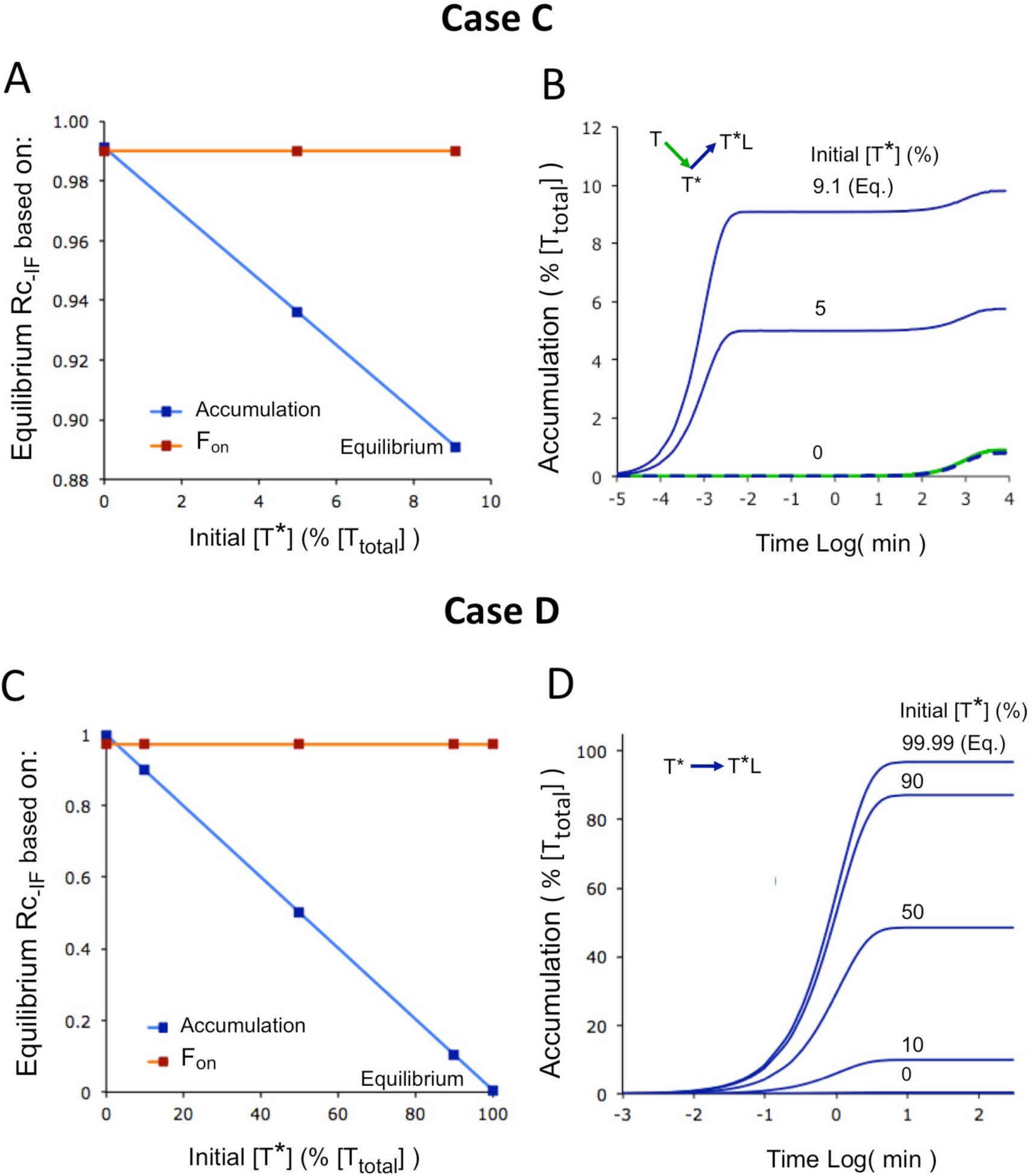
Impact of the initial [T*] on Rc_-IF_ at equilibrium binding. A) Impact of the initial [T*] on F_on_- and accumulation- based equilibrium- Rc_-IF_ values for case C. [T*] ranges from 0 till the value that corresponds to the initial [T] – [T*] equilibrium. Same [L] as in Figure 5. B) Impact of the initial [T*] on the time-wise accumulation of the target species of interest via the indicated microscopic steps (arrows) for case C. C) Impact of the initial [T*] on F_on_- and accumulation- based equilibrium- Rc_-IF_ values for case D. Same [L] as in Figure 6. D) Impact of the initial [T*] on the time-wise accumulation of [T*L] via the CS-binding step (arrow) for case D.

Comparing the microscopic step- based accumulation plots for case C in Figure 7B reveals that the early portion of the CS-binding step to [T*L] increases in par with the initial [T*]. This finding explains the positive link between the initial [T*] and the decline in accumulation- based Rc_-IF_ in Figure 7A. The similar profiles in Figure 7D indicate that such link can also be made for case D. By contrast, however, comparing Figures 6B and 6C already indicates that the CS-binding step now lags behind the IF-binding- and isomerization steps for case D when starting from an initial [T] – [T*] equilibrium. This inverted succession is even more clearly shown when starting with a lower initial [T*] (i.e. 50 % of [T_tot_]) in Supplementary Figure S8. The difference between cases C and D suggests that the above link does not necessarily require extremely fast CS-binding provided that T*-to-T retro- isomerization is highly unfavorable.

Taken together, the present examples provide further evidence for the link between a high initial [T*] and dissimilar F_on_- and accumulation- based equilibrium Rc_-IF_ values.

## Discussion

Distinct binding flux- based/related approaches have already been proposed to estimate the relative contribution of the IF- and CS pathways to the ultimate T*L complex within a thermodynamic cycle. While most researchers adopted the initial macroscopic F_-on_- based approach [22], an alternative accumulation- based approach is slowly getting momentum as well [29–31]. Both approaches provided quite comparable Rc_-IF_ – [L] relationships for most of the already reported cases but it is of note that the accumulation- based Rc_-IF_ values were about 10 % lower in (the Supplementary section of) Ordabayev et al. [29]. Based on the kinetic data for four reported cases in the literature, the present simulations show that this divergence is related to the presence of sizable [T*] in the absence of ligand. It is striking that most of those cases only partially comply with the classical paradigms for IF- and CS-binding (Supplementary Table S1). Nevertheless, this variability helped us gaining better insight into the intricate functioning of IF – CS based thermodynamic cycles. Alike previous reports [30,47], the present extended set of binding flux- based approaches shed light on the ability of both pathways to act in unison rather than being mutually exclusive. Moreover, they reveal that different approaches for calculating equilibrium- Rc_-IF_ values [22,28,29] may produce different outcomes.

The binding flux- concept is already pretty ancient [50] and the differential equations that describe how ligand binding evolves with time (Supplementary Table S2) are actually based on such fluxes. Those equations were also used for elaborating algebraic equations for kinetic parameters like pseudo first-order association rate constants, k_obs_, [9,18,23, 51,52] and for describing the competition between two one-step binding ligands [53]. Yet, they are often highly complex and, as such, not easy to decode in terms of the intimate aspects of the binding mechanism in question. Such knowledge gap can be at least partly remedied by making use of an extended range of binding flux- related approaches. Indeed, such additional tools offer alternative and visually more attractive understanding of the functioning of complex binding mechanisms such as the present thermodynamic cycle [30,31,47]. In this respect, we here illustrate that IF and CS can also act in succession in a single association experiment (Figure 6C). Along with previous observations [30,47], this finding illustrates that the actions of IF and CS can be appreciably more interwoven than traditionally suspected.

It is of note that several binding flux- based approaches offer a detailed comparison of how individual steps and pathways are able to progress with time till global equilibrium. This can already be achieved by directly comparing the progression of the microscopic- and macroscopic fluxes; i.e. all the connected forward- and reverse fluxes have become equal when global equilibrium is attained (Figures 5F and 6F). Such progression can be even more conveniently expressed in terms of their ratios; i.e. all those should equal unity (Supplementary Figure S7). Also, the accumulation of each target species via its connected steps should have come to a standstill (Figures 5C and D, 6B to D).

The accumulation- based Rc_-IF_ calculations also permit a comparison of how much the microscopic IF-isomerization- and the CS-binding steps contributed to the accumulation of T*L till any moment of interest (Table 1). It is of note that the so-obtained equilibrium-Rc_-IF_ values are very similar to those that are provided by the F_on_- approach when the initial [T*] is very low (Figure 2 and 7). However, the accumulation- based approach provides lower values at equilibrium when the initial [T*] becomes more pronounced. Regardless thereof, the accumulation- based values increase also slower for cases A to C at high [L] (Figure 5E and Supplementary Figure S6).

Those different outcomes draw attention to the fact that both approaches rely on distinct principles. First, F_on_ has to originate from the ground state (i.e. T) by definition [22] whereas the accumulation- based approach takes also account of potential contributions of any [T*] and [TL] that is already initially present. Second, while the F_on_- based approach compares fluxes at a given time point only, the accumulation- based approach acts as an archivist that takes account of early events like the fast binding of the ligand to pre-existing [T*] (Figure 5C and D). Third, the accumulation- based approach takes account of the reversible nature of all the involved microscopic steps (i.e. since it is based on the difference between forward- and reverse fluxes) whereas the F_on_- based approach only compares forward fluxes. In this respect, it is also interesting to mention that Sekhar et al. [28] did recently opt for a hybrid model by still leaning on forward fluxes but by dismissing the CS-isomerization step. Quite unexpectedly, the so-obtained Rc_-IF_ values do not vary with [L] (Supplementary Figure S9).

Case D represents an extreme situation for which the accumulation- and F_on_- based equilibrium Rc_-IF_ values diverge completely at high [L] (Figure 4C). This means that its IF pathway is able to largely dominate according to F_-on_- based- calculations despite of the fact that it only minimally contributes to the accumulation of [T*L] (Figure 6 B to D). This aberration raises questions about the pertinence of those approaches in general. Interestingly, this issue also reminds of earlier debates about how receptors are activated by an agonist (more detailed information in Supplementary Section and Figure S10). In short, the “rate theory” by Paton [54,55] assumes that each association event produces “one quantum of excitation” after which the occupied receptors become inactive again. This actually corresponds to a forward binding flux. On the other hand the “occupation theory” by Clark [56] and subsequent upgrades thereof [57–62] assumes that the receptors can be active as long as an agonist is bound. It is appealing to associate the present F_on_- based approach to the rate theory and the accumulation- based approach to the occupation theory. Indeed, while the F_on_- based approach is based on how fast the forward- as well as reverse (macroscopic) T – T*L transitions take place, the accumulation- based approach compares the contribution of IF and CS in terms of the actual target occupancies [62,63].

It is of note that the occupation theory (and in its wake, the accumulation- based approach) is largely ‘inbred’ in the pharmacological community. This state of affairs owes to a number of observations and considerations. First, several studies have shown that the efficacy agonists is positively correlated to their residence time at several G-protein-coupled receptors [64–66]. These results are not compatible with the rate theory, which requires fast dissociation (and thus a faster succession of association events) for achieving high efficacy. Second, the occurrence of constitutive receptor activity (i.e. receptor activity even in the absence of agonist) and inverse agonism (i.e. the ability of some antagonists to reduce this activity) 15,62,67 is not compatible with the “rate theory” either. Third, such as already mentioned in the introduction, the clinical benefit of a long residence time of drug-target complexes is now overwhelmingly associated with their clinical benefit. Fourth, ligand-target interactions are very often classified as either IF or CS based on the shape of their pseudo-first order association rate constant for ligand binding, k_obs,_ versus [L] plots [9,18,23,5,52].

The above considerations refer to situations that obey the “detailed balance rule” [24]. Such as for the present- and related cases, this rule allows genuine equilibrium (i.e. situation in where the forward and reverse fluxes of the all the pathways and their individual steps cancel out) to be attained. While those considerations apply to reversible drug- receptor binding and allosteric modulation, they are unlikely to do so for cases that involve enzyme activity or ion transport. In this respect, it is of note that enzyme action has already been represented almost five decades ago by a turning wheel- model [24,68]. This model is quite similar to the present IF – CS- based cycle, except that the T*L-to-T* conversion is now governed by the sum of k_4-CS_ (for the dissociation of the naïve ligand from T*L) and the rate constant of the chemical reaction (for product release). This modification was found to produce a constant clockwise- turning non-zero “net flux” within the cycle. Rather than genuine equilibrium, such cycle can thus only attain a non-equilibrium steady state in where the microscopic IF-forward- and CS-reverse fluxes exceed their respective counterparts by an equal amount.

In conclusion, ligand-target binding data are routinely fitted to more or less complex algebraic equations without even taking notice that those originate from binding flux-based differential equations. Historically, such fluxes have only attracted little attention until Hammes et al. [22] promoted the use of macroscopic forward fluxes for comparing the respective contribution of IF- and CS pathways within a thermodynamic cycle at equilibrium. Alike past work [30,31,47], the present study illustrates the pertinence of adopting even more binding flux- based approaches for better understanding the intricate functioning of complex binding mechanisms, not only for equilibrium- but also for often neglected but physiologically more relevant non-equilibrium situations [69].

## Statements

### Ethics approval, Patient consent, Permission to reproduce, Clinical trials

1. The research work does not involve the use of animal or human subjects.
2. The article is the authors’ own original work and has not been previously published, nor being considered for publication elsewhere.
3. The article reflects the authors’ own research and analysis in a truthful and complete manner.
4. The article properly credits the meaningful contributions of the co-authors.
5. Prior and existing research known by the authors is appropriately referred to.
6. All sources used are properly disclosed.

### Data Availability

No data have been shared.

### Supporting Information section- sections

S1. Microscopic rate constants for the presented Cases. (Table S1)

S2. Target concentrations at the onset and differential equations (Table S2). S3. From target concentrations to F_on_- based equilibrium R_c-IF_. (Figure S3) S4. [TL]- peak increases with [L]: e.g. Case A. (Figure S4)

S5. Ligand binding versus target activity (Figure S5)

S6. Attainment of half- maximal Rc_-IF_ for cases A and B (Figure S6)

S7. Forward/reverse flux ratios for case C. (Figure S7)

S8. Ligand association and contribution of the microscopic steps for case D (Figure S8) S9. Omission of the CS-isomerization step (Figure S9).

S10. ”Rate theory” versus “occupation theory” for receptor activation (Figure S10).

## Legend to Table 1

### Approaches for comparing the contribution of IF and CS within a thermodynamic cycle

Microscopic fluxes correspond to the product of the concentration of the starting target species and either with the associated first order rate constant or second-order rate constant and [L]. The macroscopic F_on_ for each pathway refers to the overall rate by which T converts into T*L and is related to constituent microscopic fluxes by the equations provided by Hammes et al. [22] (here only shown for F_on_). Note that, at equilibrium, F_on_ can also be obtained by the product of [T] and first-order “conditional rate” such as reported [24].
Ordabayev et al. [29] introduced an alternative [T*L]- accumulation based approach for calculating such Rc_-IF_ values. The accumulation of T*L till time t’ via each pathway can be obtained by integrating the fluxes of the last microscopic step of each pathway such as shown (please see section 2.2 for more information).
Calculation of the relative contribution of the IF pathway to [T*L] (i.e. Rc_-IF_) is based on comparing the F_on_- and [T*L]-accumulation (Acc) values of both pathways such as shown.

## Abbreviations

[ ]: Ligand or target concentrations.
F: binding flux (F_1_, F_3_ : Microscopic forward fluxes for each single reversible step of a pathway ; F_2_, F_4_ : Microscopic reverse fluxes for such steps; F_on_ : Macroscopic forward flux of a single pathway; F_off_ : Macroscopic reverse flux of such pathway (_CS-_ and _IF-_ appending’s designate the pathway).
K_D_: Thermodynamic equilibrium dissociation constant.
k_obs_: pseudo-first order association rate constant for ligand binding (also referred to as the rate, or eigenvalue, of relaxation to equilibrium).
L: Ligand (i.e. a drug or protein that binds to a target of interest)
Rc_-IF_: Relative contribution (or importance) of the IF pathway to the T*L complex. T, T* : Unbound target in the ground state and in the “activated” state, respectively.
TL, T*L: Ligand-bound target in the ground state and in the “activated” state, respectively.
[T_tot_]: Total target concentration

## References

1) Swinney DC. Biochemical mechanisms of drug action, what does it take for success? Nat Rev Drug Discov. 2004; 3:801–808. doi: 10.1038/nrd1500. PMID: 15340390

2) Copeland RA, Pompliano DL, Meek TD. Drug-target residence time and its implications for lead optimization. Nat Rev Drug Disc. 2006; 5:730–739 doi: 10.1038/nrd2082. PMID: 16888652

3) Garvey EP. Structural mechanism of slow-onset, two-step enzyme inhibition. Curr Chem Biol. 2010; 4:64–73. doi: 10.2174/187231310790226215.

4) Dror RO, Pan AC, Arlowa DH, Borhania DW, Maragakisa P, Shana Y et al. Pathway and mechanism of drug binding to G-protein-coupled receptors. Proc Natl Acad Sci USA. 2011; 108:13118–13123. doi: 10.1073/pnas.1110499108. PMID: 21778406

5) Weis WI, Kobilka BK. The Molecular Basis of G protein–coupled receptor activation Annu Rev Biochem. 2018; 87:897–919. doi: 10.1146/annurev-biochem-060614-033910. PMID: 29925258

6) De Paula VS, Jude KM, Nerli S, Glassman CR, Garcia KC, Sgourakis NG. Interleukin-2 druggability is modulated by global conformational transitions controlled by a helical capping switch. Proc Natl Acad Sci USA. 2020; 117:7183–7192. doi: 10.1073/pnas.2000419117. PMID: 32184322

7) del Castillo J, Katz B. Interaction at end-plate receptors between different choline derivatives. Proc R Soc Lond B Biol Sci. 1957; 146:369–38. doi: 10.1098/rspb.1957.0018. PMID: 13431862

8) Koshland DE. The applicaion and usefulness of the ratio kcat/KM. Bioorg Chem. 2002; 30:2011–2013. doi: 10.1006/bioo.2002.1246. PMID: 12406705

9) Strickland S, Palmer G, Masset V. Determination of dissociation constants and specific rate constants of enzyme-substrate (or protein-ligand) interactions from rapid reaction kinetic data. J Biol Chem. 1975; 250:4048–4052. doi: https://doi.org/10.1016/S0021-9258(19)41384-7 PMID: 1126943

10) Tummino PJ, Copeland RA. Residence time of receptor-ligand complexes and its effect on biological function. Biochemistry. 2008; 47:5481–5492. doi: 10.1021/bi8002023. PMID: 18412369.

11) Du X, Li Y, Xia Y-L, Ai S-M, Liang J, Sang P et al. Insights into protein-ligand interactions: mechanisms, models, and methods. Int J Mol Sci. 2016; 17:144–177. doi: 10.3390/ijms17020144. PMID: 26821017

12) Monod J, Wyman J, Changeux JP. On nature of allosteric transitions - a plausible mmodel. J Mol Biol. 1965; 12:88–118. doi: 10.1016/s0022-2836(65)80285-6. PMID: 14343300

13) Burgen AS. Conformational changes and drug action. Fed Proc. 1981; 40:2723-2728. PMID: 7297703

14) Ma B, Kumar S, Tsai CJ, Nussinov R. Folding funnels and binding mechanisms. Protein Eng. 1999; 12:713-720. doi: 10.1093/protein/12.9.713. PMID: 10506280

15) Changeux JP, Edelstein S. Conformational selection or induced fit? 50 years of debate resolved. F1000 Biol Rep. 2011; 3:19. doi: 10.3410/B3-19. PMID: 21941598.

16) Daniels KG, Tonthat NK, McClure DR, Chang YC, Liu X, Schumacher MA, et al. Ligand concentration regulates the pathways of coupled protein folding and binding. J Am Chem Soc. 2014; 136:822–825. doi: 10.1021/ja4086726. PMID: 24364358

17) Greives N, Zhou H-X. Both protein dynamics and ligand concentration can shift the binding mechanism between conformational selection and induced fit. Proc Nat Acad USA. 2014; 111:10197–10202. doi: 10.1073/pnas.1407545111. PMID: 2498214

18) Meyer-Almes FJ. Discrimination between conformational selection and induced fit protein-ligand binding using Integrated Global Fit analysis. Eur Biophys J. 2016; 45:245–257. doi: 10.1007/s00249-015-1090-1. PMID: 26538331

19) Copeland RA. The dynamics of drug-target interactions: drug-target residence time and its impact on efficacy and safety. Expert Opin Drug Discov. 2010; 5:305–310. doi: 10.1517/17460441003677725. PMID: 22823083

20) Copeland RA. Conformational adaptation in drug-target interactions and residence time. Future Med Chem. 2011; 3:1491–1501. doi: 10.4155/fmc.11.112. PMID: 21882942

21) Boehr DD, Nussinov R, Wright PE. The role of dynamic conformational ensembles in biomolecular recognition. Nat Chem Biol. 2009; 5:789–796. doi: 10.1038/nchembio.232. PMID: 19841628

22) Hammes GG, Chang YC, Oas TG. Conformational selection or induced fit: a flux description of reaction mechanism. Proc Natl Acad Sci USA. 2009; 106:13737–13741. doi: 10.1073/pnas.0907195106. PMID: 19666553.

23) Vogt AD, Di Cera E. Conformational selection or induced fit? A critical appraisal of the kinetic mechanism. Biochemistry. 2012; 51:5894–5902. doi: 10.1021/bi3006913. PMID: 22775458

24) Michel D. Conformational selection or induced fit? New insights from old principles. Biochimie. 2016; 128–129:48–54. doi: 10.1016/j.biochi.2016.06.012. PMID: 27344613

25) Li D, Ji B. Protein conformational transitions coupling with ligand interactions: Simulations from molecules to medicine. Medicine in Novel Technology and Devices. 2019; 3:100026. doi: https://doi.org/10.1016/j.medntd.2019.100026. PMID: 27344613

26) Galburt EA, Rammohan J. Kinetic signature for parallel pathways: conformational selection and induced fit. Links and disconnects between observed relaxation rates and fractional equilibrium flux under pseudo-first-order conditions. Biochemistry. 2016; 55:7014–7022. doi: 10.1021/acs.biochem.6b00914. PMID: 27992996

27) Zhou G, Pantelopoulos GA, Mukherrjee S, Voelz VA. Bridging microscopic and macroscopic mechanisms of p53-MDM2 binding with kinetic network models. Biophys J. 2017; 113: 785–793. doi: 10.1016/j.bpj.2017.07.009. PMID: 28834715

28) Sekhar A, Velyvis A, Zoltsman G, Rosenzweig G, Bouvignies G, Kay LE. Conserved conformational selection mechanism of HP70 chaperone-substrate interactions. Elife. 2018; 7:e32764. doi: 10.7554/eLife.32764. PMID: 29460778

29) Ordabayev YA, Nguyen B, Kozlov AG, Jia H, Lohman TM. UvrD helicase activation by MutL involves rotation of its 2B sub-domain. Proc. Natl. Acad. Sci. USA. 2019; 116: 16320–16325. doi: 10.1073/pnas.1905513116. PMID: 31363055

30) Vauquelin G, Maes D, Swinney DC. Fluxes for unraveling complex binding mechanisms. Trends Pharmacol Sci. 2020; 41:923–932. doi: 10.1016/j.tips.2020.10.003 PMID: 33153779

31) Vauquelin G, Maes D. Induced fit versus conformational selection: from rate constants to fluxes… and back to rate constants. Fund Res Perspect. 2021; 9:e00874. doi: 10.1002/prp2.847. PMID: 34459109

32) Ye L, Van Eps E, Zimmer M, Ernst OP, Scott Prosser R. Activation of the _A2A_ adenosine G-protein-coupled receptor by conformational selection. Nature. 2016; 533:265–268. doi: 10.1038/nature17668. PMID: 27144352

33) Her C, Thompson AR, Karim CB, Thomas DD. Structural dynamics of calmodulin-ryanodine receptor interactions: electron paramagnetic resonance using stereospecific spin labels. Scientific Reports. 2018; 8:10681 doi: 10.1038/s41598-018-29064-8. PMID: 3001309

34) Hu W, Wang H, Hou Y, Hao Y, Liu D. Trimethylsilyl reporter groups for NMR studies of conformational changes in G protein-coupled receptors. FEBS Lett. 2019; 593,1113–1121. doi: 10.1002/1873-3468.13382. PMID: 30953343

35) Wu F-J, Williams LM, Abdul-Ridha A, Gunatilaka A, Vaid TM., Kocan M et al. Probing the correlation between ligand efficacy and conformational diversity at the α_1A_-adrenoreceptor reveals allosteric coupling of its microswitches. J Biol Chem. 2020; 295: 7404–7417. doi: 10.1074/jbc.RA120.012842. PMID: 32303636

36) Shimada I, Ueda T, Kofuku Y, Eddy MT, Wüthrich K. GPCR drug discovery: integrating solution NMR data with crystal and cryo-EM structures. Nat Rev Drug Discov. 2020;18: 59–82. doi: 10.1038/nrd.2018.180. PMID: 30410121

37) Roither B, Oostenbrink C, Pfeiler G, Koelbl H, Schreiner W. Pembrolizumab induces an unexpected conformational change in the CC’-loop of PD-1. Cancers. 2021; 13:5. doi: 10.1038/nrd.2018.180. PMID: 33375020

38) Fuxreiter M. Classifying the binding modes of disordered proteins. Int J Mol Sci. 2020; 21:8615. doi: 10.3390/ijms21228615. PMID: 33207556

39) Neubig R, Spedding M, Kenakin T, Christopoulos A. International Union of Pharmacology Committee on Receptor Nomenclature and Drug Classification. XXXVIII. Update on terms and symbols in quantitative pharmacology. Pharmacol Rev. 2003; 55:597–606. doi: 10.1124/pr.55.4.4. PMID: 14657418

40) Vauquelin G, Morsing P, Fierens FLP, De Backer JP, Vanderheyden PML. A two-state receptor model for the interaction between angiotensin II AT_1_ receptors and their non-peptide antagonists. Biochem Pharmacol. 2001; 61:277–284. doi: 10.1016/s0006-2952(00)00546-3. PMID: 11172731

41) Vauquelin G, Charlton S. Exploring avidity: understanding the potential gains in functional affinity and target residence time of bivalent and heterobivalent ligands. Brit J Pharmacol. 2013; 168:1771–1785. doi: 10.1111/bph.12106. PMID: 23330947

42) Malygin EG, Hattman SA. probabilistic approach to compact steady-state kinetic equations for enzymic reactions. J Theor Biol. 2006; 242:627–633. doi: 10.1016/j.jtbi.2006.03.022. PMID: 16697416

43) Chakraborty P, Di Cera P. Induced fit is a special case of conformational selection. Biochemistry. 2017; 56:2853–2859. doi: 10.1021/acs.biochem.7b00340. PMID: 28494585

44) Vauquelin G. Distinct *in vivo* target occupancy by bivalent- and induced-fit-like binding drugs. Br J Pharmacol. 2017; 173:1268–1285 doi: 10.1111/bph.13989. PMID: 28838028

45) Dahl G, Akerud T. Pharmacokinetics and the drug-target residence time concept. Drug Discov Today. 2013; 18:697–707. doi: 10.1016/j.drudis.2013.02.010. PMID: 23500610

46) Vauquelin G, Van Liefde I, Swinney D. On the different experimental manifestations of two-state “induced fit” binding of drugs to their cellular targets. Br J Pharmacol. 2016; 173:1268–1285. doi: 10.1111/bph.13445. PMID: 26808227

47) Vauquelin G, Maes D. Competition in drug binding and … the race to equilibrium. Fund Clin Pharmacol. 2023; 37:147–157. doi : 10.1111/fcp.12824. PMID: 35981720

48) Swinney DC, Beavis P, Chuang K-T, Zheng Y, Lee I, Gee P et al. A study of the molecular mechanism of binding kinetics and long residence times of human CCR_5_ receptor small molecule allosteric ligands. Br J Pharmacol. 2014; 171:3364–3375. doi: 10.1111/bph.12683. PMID: 24628038

49) Vauquelin G, Van Liefde I, Swinney D. On the different experimental manifestations of two-state “induced fit” binding of drugs to their cellular targets. Br J Pharmacol. 2016; 173:1268–1285. doi: 10.1111/bph.13445. PMID: 26808227

50) Hill TL, Chen Y-D. Stochastics of cycle completions (Fluxes) in biochemical kinetic diagrams. Proc Natl Acad Sci USA. 1975; 72: 291–1295. doi: 10.1073/pnas.72.4.1291. PMID: 1055403

51) Gianni S, Dogan J, Jemth P. Distinguishing induced fit from conformational selection. Biophys Chem. 2014; 189:33–39. doi: 10.1016/j.bpc.2014.03.003. PMID: 24747333

52) Paul F, Weikl TR. How to distinguish conformational selection and induced fit based on chemical relaxation rates. PLoS Comput Biol. 2016; 12: e1005067. doi: 10.1371/journal.pcbi.1005067. PMID: 27636092

53) Motulsky HJ, Mahan LC. The kinetics of competitive radioligand binding predicted by the law of mass action. Mol Pharmacol. 1984; 25:1–9. PMID: 6708928

54) Paton WDM. A theory of drug action based on rate of drug-receptor combination. Proc R Soc Lond B Biol Sci. 1961; 154:21–69. URL: https://www.jstor.org/stable/90247

55) Paton WDM. Kinetic theories of drug action with special reference to the acetyccholine group of agonists and antagonists. Ann NY Acad Sci. 1967;144:869–881. doi: 10.1111/j.1749-6632.1967.tb53816.x

56) Clark AJ. General Pharmacology: in: ‘Heffner’s Handbuch Der Experimentellen Pharmacologie Ergänzungsband, Band 4. Springer-Verlag: Berlin (1937).

57) Ariens EJ. Affinity and intrinsic activity in the theory of competitive inhibition. 1. Problems and theory. Arch Int Pharmacodyn Ther. 1954; 99:32–49. PMID: 13229418

58) Stephenson RP. A modification of receptor theory Br J Pharmacol. 1956; 11:379–393. doi: 10.1111/j.1476-5381.1956.tb00006.x. PMID: 13383117

59) Furchgott RF. The use of β-haloalkylamines in the differentiation of receptors and in the determination of dissociation constants of receptor-agonist complexes. In: Harper NJ, Simmonds AB (eds). Advances in Drug Research., Vol. 3. Academic Press: New York, 1966; pp. 21-55.

60) Leff P. The two-state model of receptor activation. Trends Pharmacol Sci. 1995; 16:89–97. doi: 10.1016/s0165-6147(00)88989-0. PMID: 7540781

61) Colquhoun D. Binding, gating, affinity and efficacy: the interpretation of structure-activity relationships for agonists and of the effects of mutating receptors. Br J Pharmacol. 1998; 125:924–947. doi: 10.1038/sj.bjp.0702164. PMID: 9846630

62) Kenakin T. New concepts in pharmacological efficacy at 7TM receptors: IUPHAR Review 2. Br J Pharmacol. 2012; 168:554–575. doi: 10.1111/j.1476-5381.2012.02223.x. PMID: 22994528

63) Bosshard HR. Molecular recognition by induced fit: how fit is the concept? News Physiol Sci. 2001; 16:171–173. doi: 10.1152/physiologyonline.2001.16.4.171. PMID: 11479367

64) Sykes DA, Dowling MR, Charlton SJ. Exploring the mechanism of agonist efficacy: A relationship between efficacy and agonist dissociation rate at the muscarinic M_3_ receptor. Mol Pharmacol. 2009; 76:543–551. doi: 10.1124/mol.108.054452. PMID: 19498041

65) Guo D, Mulder-Krieger T, IJzerman AP, Heitman LH. Functional efficacy of adenosine A_2_A receptor agonists is positively correlated to their receptor residence time. Br J Pharmacol. 2012; 166:1846–1859. doi: 10.1111/j.1476-5381.2012.01897.x. PMID: 22324512

66) Rosethorne EM, Bradley ME, Gherbi K, Sykes DA, Sattikar A, Wright JD et al. Long receptor residence time of C26 contributes to super agonist activity at the human ß_2_ adrenoceptor. Mol Pharmacol. 2016; 89:467–475. doi: 10.1124/mol.115.101253. PMID: 26772612

67) Lefkowitz RJ, Cotecchia S, Samama P, Costa T. Constitutive activity of receptors coupled to guanine nucleotide regulatory proteins. Trends Pharmacol Sci. 1993;14:303–307. doi: 10.1016/0165-6147(93)90048-O. PMID: 8249148

68) Wyman J. The turning wheel: a study in steady states, Proc Natl Acad Sci USA. 1975; 72: 3983–3987. doi: 10.1073/pnas.72.10.3983. PMID: 1060079

69) Copeland RA. The drug-target residence time model: a 10-year retrospective. Nat Rev Dug Discov. 2016; 15:87–95. doi: 10.1038/nrd.2015.18. PMID: 26678621

