## Supplementary tables and figures for "Competition between induced fit- and conformational selection binding: how different flux-based approaches help clarifying their interplay"

Tables: 2

Figures: 8

**Table of Contents** Page

S1. Microscopic rate constants for the presented Cases. (Table S1) 2

S2. Target concentrations at the onset and differential equations (Table S2). 3

S3. From target concentrations to $F_{on}$- based equilibrium $R_{c\text{-}IF}$. (Figure S3) 3

S4. [TL]- peak increases with [L]: e.g. Case A. (Figure S4) 4

S5. Ligand binding versus target activity (Figure S5) 4

S6. Attainment of half- maximal $Rc_{\text{-}IF}$ for cases A and B (Figure S6) 5

S7. Forward/reverse flux ratios for case C. (Figure S7) 5

S8. Ligand association and contribution of the microscopic steps for case D (Figure S8) 6

S9. Omission of the CS-isomerization step (Figure S9). 7

S10. ”Rate theory” versus “occupation theory” for receptor activation (Figure S10). 8

Tables 10

Figures 12

References 18

**S1. Microscopic rate constants for the presented Cases. (Table S1)**

The microscopic rate constants of the presented cases are given in Panel A. Panel B shows relevant ratios thereof and examines their compliance with the classical paradigms for IF and CS binding. Since the original constants for case B were all extremely high, they were presently divided by $10^6$ for the sake of legibility.

**Specific comments.**

The "detailed balance rule" [4] stipulates that the difference in Gibbs free energy between T and T*L needs to be the same for the IF- and CS pathways of a thermodynamic cycle. This implies that the thermodynamic $K_D$'s (i.e. the $k_2.k_4/k_1.k_3$- ratios) of those two pathways have to be equal as well. The presented cases obey this rule. This implies that, when only approximate rate constants are provided in the literature, one of them has to be slightly adjusted.

$k_{3\text{-IF}}/k_{2\text{-IF}}$ ratios : Genuine IF binding should go along with a fast binding step, followed by a slower isomerization step whereas "bivalent-like" IF binding should go along with a slow binding step, followed by a faster isomerization step [5]. Accordingly, $k_{3\text{-IF}}/k_{2\text{-IF}}$ ratio should be < 1 for genuine IF binding and > 1 for "bivalent-like" IF binding [6,7]. In this respect, it is of note that the $k_{3\text{-IF}}/k_{2\text{-IF}}$ ratios are equal to- or close to 1 for cases A and D, i.e. two examples that were provided by Hammes et al. [1]. As such, they represent hybrid situations.

$k_{4\text{-IF}}/k_{3\text{-IF}}$ ratios : Such as shown in Figures 2 and 3 of the article, this ratio should be < 1 to allow [T*L] to represent nearly all (cases A and D), or at least the majority (case C) of the bound targets at equilibrium in the presence of ligand. This condition is clearly not met for case B.

$k_{2\text{-CS}}/k_{1\text{-CS}}$ ratios : This ratio defines the amount of "active" receptors, [T*], when the cycle is in equilibrium in the absence of ligand, i.e. at the onset of the present simulations (except for Figure S8). This ratio should be >1 when, according to the classic definition of CS binding, T* represents a thermodynamically unstable high- energy conformation so that [T*] only constitutes a minute fraction of the total target population. Yet, the rate constants that were provided by Hammes et al. [1] for their second example (case D) yield a radically opposite ratio. Such as shown in Figure 3 of the article, this deviation explains why [T*] nearly represents almost all of the unbound targets at the initial equilibrium.

$k_{4\text{-}CS}/k_{1\text{-}CS}$ ratios : $k_{obs}$ Versus [L] plots are still widely used to differentiate IF- form CS binding. Those plots were initially thought to increase hyperbolically for IF and to decrease for CS [5,8]. This distinction was based on the then prevailing premise that the conformational change of those pathways is much slower than the actual binding process. For CS, this requires the $k_{4\text{-}CS}/k_{1\text{-}CS}$ ratio to exceed unity [9,10]. However, $k_{obs}$ versus [L] plots may be hyperbolically as well if the CS-isomerization step proceeds faster than its binding step, i.e. when $k_{4\text{-}CS}/k_{1\text{-}CS} < 1$. As such, case A represents a hybrid situation whose $k_{4\text{-}CS}/k_{1\text{-}CS}$ ratio equals 1 whereas case D represents a situation with a very fast CS-isomerization step.

#### S2. Target concentrations at the onset and differential equations (Table S2).

Differential equations 1 to 4 account for the time- wise accumulation (i,e. evolution of the concentration) of the different free- and ligand-bound target species shown on top of Table S1. Those equations have to be integrated in parallel such as stipulated in Section 2.2 of the article.
Differential equations 5 to 9 account for the steps that provide a positive contribution to the accumulation of the target species of interest. They need to be integrated alongside with equations 1 to 4.

**Target concentrations for the onset (i.e. t = 0) at equilibrium are:**

$[T^*]$ (in % of $[T_{total}]$) $= 100/(1 + k_{2\text{-}CS}/k_{1\text{-}CS})$ , $[T] = 100 - [T^*]$ , $[TL] = [T^*L] = 0$

Ligands are assumed to be in large excess over the targets, so that their concentration in the bulk of the aqueous phase remains constant over time.

#### S3. From target concentrations to $F_{on}$- based equilibrium $Rc_{\text{-}IF}$ (Figure S3)

Figure S3 provides a step-by step account of how the macroscopic $F_{on}$- based $Rc_{\text{-}IF}$ values can be calculated for case A. Please note that the calculation of $Rc_{\text{-}IF}$ requires prior calculation of of Fon-IF and Fon-CS, that those require prior calculation of F1-IF, F3-IF, F1-CS and F3-CS and that those require prior calculation of the time-wise evolution of [T], [TL], [T*] and

[T*L]. The present panels show the time-wise evolution of pertinent parameters in chronological order.

First step (Panel A). The time-wise evolution of [T], [TL], [T*] and [T*L] can be calculated by integrating equations 1 to 4 in Section S2 with the rate constants provided in Section S1, [L] (= 10 μM) and the initial concentration of the different target species ([T] = 98.03 %, [TL] = 0 %, [T*] = 1.96 % and [T*L] = 0 % of [Ttotal] (for the cycle in initial equilibrium) as input. Target concentrations that are important for the calculation of F1-IF, F3-IF are shown at the left side and those for the calculation of F1-CS and F3-CS are shown at the right side. Inserts are magnifications.

Second step (Panel B). The time-wise evolution of the individual microscopic and macroscopic forward fluxes can then be calculated by the provided equations For IF (left side) and for CS (right side). Those macroscopic values are largely dictated by the lowest of the microscopic fluxes, here mainly F3-IF for Fon-IF and F1-CS for Fon-CS. Please note that Fon is only half of F1 and F3 when their values are equal. Panel C compares the time-wise evolution of Fon-IF and Fon-CS.

Third step (Panel D). The time- wise evolution of the relative contribution of Rc-IF, can then be calculated based on the macroscopic forward fluxes according to the equation shown.

**S4. [TL]- overshoot increases with [L]: e.g. Case A. (Figure S4)**

Figure S4 compares the time-wise changes in the concentration of the intermediary- and the ultimate T*L species for the same [L] as in Figure S3 (Panel A) and for a 10- fold higher [L] (panel B). The transient [TL]-overshoot is clearly more pronounced at high [L].

**S5. Ligand binding vs. target activity (Figure S5)**

Figure S5 compares the time- wise evolution of the “observable” binding of L in e.g. radioligand association experiments (i.e. [TL] + [T*L]) with the “observable” target activity (i.e. [T*] + [T*L]) for the four cases that are presented in the article. Those simulations are based on the premise that [TL] is sufficiently long lasting to be observable in the involved experimental settings such as radioligand binding. Also, please note that T* and T*L are generally considered to represent "active" target species with respect to e.g. G- protein-coupled receptors [11-13]. Although they are presently considered to be equally active, this might not always be the case [13,14].

The concentrations of the individual target species are taken from Figure 3 of the article. It is only for case A that ligand binding and activity evolve nearly identical. Ligand binding precedes the activity for the cases B and C and the other way round for Case D. These differences can be imputed to the [TL] overshoot for the former cases and the already excessively high [T*] for the latter case.

#### S6. Attainment of half- maximal $Rc_{-IF}$ for cases A and B (Figure S6)

Figure S6 compares the time-wise progression the $F_{on}$- and accumulation- based $Rc_{-IF}$ values till their final plateau for case A and B at high [L] (i.e. 10 µM for case A and 30 mM for case B).

Left-side panels: Black bars refer to [L] at which [T*L] is half-maximal. Such as already documented for case C in Figure 5 of the article, the $F_{on}$- based $Rc_{-IF}$ values progress here also faster than the accumulation- based counterparts.

Right-side panels: The underlying mechanisms are also the same as for Case C in the article. Indeed, the accumulation- based $Rc_{-IF}$ values reach the 0.5- mark when the IF and CS contribute equally to [T*L]: i.e. when the TL ➞ T*L transition (orange) finally catches up with the prior contribution of the CS-binding step (blue). On the other hand, the $F_{on}$- based $Rc_{-IF}$ values reach the 0.5- mark when the lowest of the microscopic fluxes of each pathway (i.e. here also $F_{3\text{-}IF}$ and $F_{1\text{-}CS}$) do cross. Only those fluxes are shown.

#### S7. Forward/reverse flux ratios for case C. (Figure S7)

Figure S7 compares the time-wise changes of the forward/reverse flux ratios for case C. Conditions are the same as for Figure 3 of the article and similar data have already been reported previously [15].

Left-side panel: In short: the initial equilibrium between [T] and [T*] explains why $F_{2\text{-}CS}/F_{1\text{-}}$

$_{CS}$ = 1 at the start. Binding of L produces a very fast conversion of T* into T*L so that the initial [T*] (and thus the $F_{2\text{-}CS}/F_{1\text{-}CS}$ ratio, green) drops quickly to nearly baseline. The therewith- associated increase of [T*L] goes along with a fast increase of the $F_{4\text{-}CS}/F_{3\text{-}CS}$ ratio (blue) till almost 1. Binding of L will also produce a fast conversion of T into TL so that the $F_{2\text{-}IF}/F_{1\text{-}IF}$ ratio (red) does rapidly increase till almost 1 as well. These early events thus indicate that both binding steps tend to reach their individual microscopic equilibrium first. The slow isomerization steps will then give rise to a late progression of the $F_{4\text{-}IF}/F_{3\text{-}IF}$- (orange) and the $F_{2\text{-}CS}/F_{1\text{-}CS}$ ratios to 1.
Right-side panel: Those isomerization steps will thus dictate the progression of the $F_{on}/F_{off}$-ratios of both pathway and also how fast the cycle attains its final equilibrium.

**S8. Ligand association and contribution of the microscopic steps for case D. (Figure S8).**

The initial [T*] now equals 50 % of [$T_{tot}$].

Panel A: The saturation binding plot shows how the concentration of each target species of case D evolves with time for [L] = 10 µM. Compared to a linear time scale, the logarithmic time scale thereof permits a straightforward comparing the processes that take place at different timeframes. First, [TL] rises very swiftly and declines only slowly afterwards. This pattern was not perceptible when dealing with an initial [T] - [T*] equilibrium in the absence of ligand such as in Figure 6A of the article. Compared to the extremely low initial [T] in the article, it presently already amounts 50 % of [$Tt_{ot}$]. This difference hints at a causal relationship between the initial [T] and the amplitude of the [TL]-overshoot. Second, the rapid full decline of the initial [T] and the subsequent slower but equal initial increase of [T*L], pleads in favor of successive intervention of the IF- binding- and isomerization steps. Third, the slower additional increase of [T*L] goes along with an equally slow full decrease of [T*]. This suggests the involvement of the CS- binding step.

Panel B: The above suggestions are confirmed by comparing the contribution of the microscopic steps to the accumulation of each target species over time (indicated with arrows). Indeed, the initial IF-binding step is followed by a slower IF-isomerization step and the even slower CS-binding step contributes equally to [T*L]. This implies that the IF-and

CS-pathways contribute equally to the accumulation of [T*L]. Moreover, Panel B also unveils a minute positive contribution of the CS- isomerization step to the accumulation of [T*]. Hence, this step now acts in the opposite way in comparison to the situation in where the initial [T] was extremely low in Figure 6 of the article. Here again, this reversal can be attributed to the presently much higher initial [T].

**S9. Ommission of the CS- isomerization step (Figure S9).**

Sekhar et al. [14] did recently opt for a hybrid situation by still leaning on forward fluxes but by dismissing the CS- isomerization step when it was too slow to be of significance during the duration of an experiment. The calculated equilibrium- $Rc_{-IF}$ thus merely relied on comparing $F_{on\text{-}IF}$ with $F_{3\text{-}CS}$ at equilibrium. Figure S9 shows the outcome of such comparison for the four cases shown in Figure 4 of the article (cases are presented in the same order). The so-obtained $Rc_{-IF}$ values (green) remain equal to the minimal $Rc_{-IF}$ for the unabridged cycle (black) at all [L].

**Mathematical considerations:**

Hammes et al. [1] referred to $F_{on\text{-}IF}$ and $F_{on\text{-}CS}$ as the macroscopic fluxes that relate the overall conversion of T into T*L. Equilibrium binding represents a special condition in where $F_{on}$ equals $F_{off}$ for each pathway. Accordingly, we can express the relative contribution of IF as:

$$Rc_{-IF} = \frac{F_{on-IF}}{F_{on-IF}+F_{on-CS}} = \frac{F_{off-IF}}{F_{off-IF}+F_{off-CS}} \qquad \text{equation 10}$$

At equilibrium, $F_{off\text{-}IF}$ and $F_{off\text{-}CS}$ can also be expressed in terms of their respective conditional rate- based equations [4,7].

$[\,T^*L\,].\,k_{2-IF}.\,k_{4-IF}\,/(k_{2-IF}+k_{3-IF})$ and $[\,T^*L\,].\,k_{2-CS}.\,k_{4-CS}\,/(k_{2-CS}+[L].\,k_{3-CS})$

Replacing $F_{on\text{-}IF}$ and $F_{on\text{-}CS}$ in equation 10 by these conditional rate- based parameters and dividing numerator and denominator by [T*L] [7] yields:

$$Rc_{-IF} = \frac{1}{1+\frac{\alpha_{F_{off}}}{k_{2-CS}+k_{3-CS}.[L]}} \quad \text{with} \quad \alpha_{F_{off}} = \frac{k_{2-CS}.k_{4-CS}.(k_{2-IF}+k_{3-IF})}{k_{2-IF}.k_{4-IF}} \qquad \text{equation 11}$$

This equation recounts the ascending shape the $F_{on}$- based $Rc_{-IF}$ versus [L] plots such as shown in Figure 4 of the article.

In contrast, Sekhar et al. [14] only compared $F_{on-IF}$ (i.e the macroscopic flux that relates the overall conversion of T into T*L via the IF- pathway) to $F_{3-CS}$ instead of $F_{on-CS}$ for the calculation of $Rc_{-IF}$. Since the microscopic forward and reverse fluxes are also equal at equilibrium, we can replace equation 1 by:

$$Rc_{-IF} = \frac{F_{off-IF}}{F_{off-IF}+F_{4-CS}} = \frac{[\,T{*}L\,].k_{2-IF}.k_{4-IF}\,/(k_{2-IF}+k_{3-IF})}{([\,T{*}L\,].k_{2-IF}.k_{4-IF}\,/(k_{2-IF}+k_{3-IF}))+[\,T{*}L\,].k_{4-CS}} \qquad \text{equation 12}$$

Dividing numerator and denominator by [T*L] yields:

$$Rc_{-IF} = \frac{k_{2-IF}.k_{4-IF}\,/(k_{2-IF}+k_{3-IF})}{(k_{2-IF}.k_{4-IF}\,/(k_{2-IF}+k_{3-IF}))+k_{4-CS}} = \frac{k_{2-CS}}{k_{2-CS}+\alpha F_{off}} \qquad \text{equation 13}$$

Please note that $R_{c-IF}$ no longer depends on [L] in equation 13 and that its value corresponds to the minimal $R_{c-IF}$- value (i.e. at [L] = 0) when comparing $F_{on-IF}$ and $F_{on-CS}$ for the unabridged cycle (i.e. Supplemental equation 6 for [7]).

#### S10. ”Rate theory” versus “occupation theory” for receptor activation (Figure S10).

Two major theories have been proposed for explaining how agonists can activate their receptors.

Panel A: On the one hand, the initial so called “rate theory” by Paton [16,17] assumed that each association-event produces “one quantum of excitation” after which the occupied receptors become inactive again. In other words, the receptors were regarded to behave as short- acting on - off switches. As such, they should display higher activity with rapid associating and dissociating agonists. At equilibrium, higher receptor activity should thus be obtained with drugs that permit more frequent association events to each single receptor molecule or, a higher number of association and dissociation events during a small time frame (Δt). This actually corresponds to a forward binding flux. Indeed, the $F_{on}$- based approach for calculating $Rc_{\text{-IF}}$ is based on comparing the number of forward- as well as reverse T – T*L- transitions during such small time frame (designated by the bidirectional arrows in the right-side panel).

Panel B: On the other hand, the "occupation theory" by Clark [18] relied on the concept that a receptor remains active as long as the agonist occupies it [17]. This theory has subsequently been upgraded in order to explain how some agonists do produce higher levels of receptor activation and response than others. This led to early popular concepts like "intrinsic activity", "efficacy" and "intrinsic efficacy" [19-21] (reviewed in [22]) as well as to more recent models that relate agonist's efficacy to its ability to stabilize an active conformation of the receptor [23,24]. Quite similarly, the accumulation- based approach for calculating $Rc_{\text{-IF}}$ values relies on comparing concentrations (right-side panel). At any time point, [T*L] is the resultant of past accumulations via IF and CS. In spite of the still occurring bidirectional T – T*L- transitions, the accumulation [T*L] has come to an end at genuine equilibrium. Please not that the accumulation of [T*L] only depends on the respective contributions of the IF- isomerization- and CS-binding steps (arrows in bold in the left-side panel).

#### Table S1

##### A) Microscopic rate constants and thermodynamic $K_D$ values.

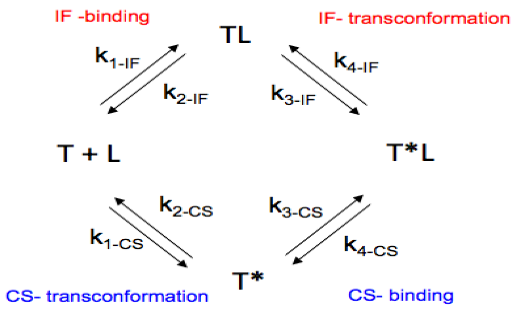


**Case A (example 1 in Hammes et al.)** [1]

| | | | | | |
|---|---|---|---|---|---|
| IF Pathway | $k_1$ ($M^{-1}.min^{-1}$) | $k_2$ ($min^{-1}$) | $k_3$ ($min^{-1}$) | $k_4$ ($min^{-1}$) | $K_D$ (M) |
| | $1\ 10^{7}$ | $1\ 10^{3}$ | $1\ 10^{3}$ | 10 | $1\ 10^{-6}$ |
| CS Pathway | $k_1$ ($min^{-1}$) | $k_2$ ($min^{-1}$) | $k_3$ ($M^{-1}.min^{-1}$) | $k_4$ ($min^{-1}$) | $K_D$ (M) |
| | 20 | $1\ 10^{3}$ | $1\ 10^{9}$ | 20 | $1\ 10^{-6}$ |

**Case B (Zhou et al.)** [2]

| | | | | | |
|---|---|---|---|---|---|
| IF Pathway | $1.8\ 10^{3}$ | 2.9 | 0.96 | 3.92 | $6.585\ 10^{-3}$ |
| CS Pathway | $1.1\ 10^{-2}$ | 10 | $4\ 10^{3}$ | $2.9\ 10^{-2}$ | $6.585\ 10^{-3}$ |

**Case C (Galburt and Rammohan.** [3]

| | | | | | |
|---|---|---|---|---|---|
| IF Pathway | $1\ 10^{8}$ | 100 | $1\ 10^{-3}$ | $1\ 10^{-4}$ | $1\ 10^{-7}$ |
| CS Pathway | $1\ 10^{-4}$ | $1\ 10^{-3}$ | $1\ 10^{8}$ | 1 | $1\ 10^{-7}$ |

**Case D (example 2 in Hammes et al.)** [1]

| | | | | | |
|---|---|---|---|---|---|
| IF Pathway | $1\ 10^{8}$ | 580 | 568 | $9\ 10^{-3}$ | $9.19\ 10^{-11}$ |
| CS Pathway | 45.34 | $5\ 10^{-3}$ | $3\ 10^{4}$ | $2.5\ 10^{-2}$ | $9.19\ 10^{-11}$ |

##### B) Compliance with the classical paradigms for IF and CS binding:

| Case: | A | B | C | D |
|---|---|---|---|---|
| IF pathway: $k_3/k_2$ | 1 | 0.33 | $1\ 10^{-5}$ | 0.98 |
| IF pathway: $k_4/k_3$ | $1\ 10^{-2}$ | 4.09 | 0.1 | $1.58\ 10^{-5}$ |
| CS pathway: $k_2/k_1$ | 50 | 910 | 10 | $1.1\ 10^{-4}$ |
| CS pathway: $k_4/k_1$ | 1 | 2.63 | $1\ 10^{4}$ | $5.51\ 10^{-4}$ |

Values in red refer to poor compliance.

#### Table S2

**A) Differential equations for how the concentration of each target species evolves with time.**

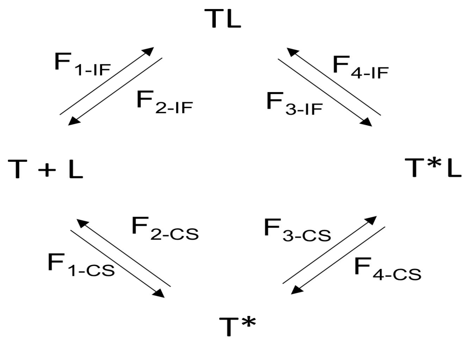


$d[T]/dt = k_{2\text{-}IF}.[TL] + k_{2\text{-}CS}.[T^*] - k_{1\text{-}IF}.[T].[L] - k_{1\text{-}CS}.[T]$ equation 1

$(= F_{2\text{-}IF} + F_{2\text{-}CS} - F_{1\text{-}IF} - F_{1\text{-}CS})$

$d[TL]/dt = k_{1\text{-}IF}.[T].[L] + k_{4\text{-}IF}.[T^*L] - k_{2\text{-}IF}.[TL] - k_{3\text{-}IF}.[TL]$ equation 2

$(= F_{1\text{-}IF} + F_{4\text{-}IF} - F_{2\text{-}IF} - F_{3\text{-}IF})$

$d[T^*]/dt = k_{1\text{-}CS}.[T] + k_{4\text{-}CS}.[T^*L] - k_{2\text{-}CS}.[T^*] - k_{3\text{-}CS}.[T^*].[L]$ equation 3

$(= F_{1\text{-}CS} + F_{4\text{-}CS} - F_{2\text{-}CS} - F_{3\text{-}CS})$

$d[T^*L]/dt) = k_{3\text{-}IF}.[TL] + k_{3\text{-}CS}.[T^*].[L] - k_{4\text{-}IF}.[T^*L] - k_{4\text{-}CS}.[T^*L]$ equation 4

$(=F_{3\text{-}IF} + F_{3\text{-}CS} - F_{4\text{-}IF} - F_{4\text{-}CS})$

**B) Differential equations for the contribution of single steps to the accumulation of a target species.**

For the rightward- pointing arrows in the figures of the article.

$d[TL]/dt = k_{1\text{-}IF}.[T].[L] - k_{2\text{-}IF}.[TL]$ $(= F_{1\text{-}IF} - F_{2\text{-}IF})$ equation 5

$d[T^*]/dt = k_{1\text{-}CS}.[T] - k_{2\text{-}CS}.[T^*]$ $(= F_{1\text{-}CS} - k_{2\text{-}CS})$ equation 6

$d[T^*L]/dt = k_{3\text{-}IF}.[TL] - k_{4\text{-}IF}.[T^*L]$ $(= F_{3\text{-}IF} - F_{4\text{-}IF})$ (via IF) equation 7

$d[T^*L]/dt = k_{3\text{-}CS}.[T^*].[L] - k_{4\text{-}CS}.[T^*L]$ $(= F_{3\text{-}CS} - F_{4\text{-}CS})$ (via CS) equation 8

For the leftward- pointing arrow in Figure 6B to D of the article: please replace Equation 6 by Equation 9.

$d[T]/dt = k_{2\text{-}CS}.[T^*] - k_{1\text{-}CS}.[T]$ $(= F_{2\text{-}CS} - F_{1\text{-}CS})$ (via CS) equation 9

**Figure S3**

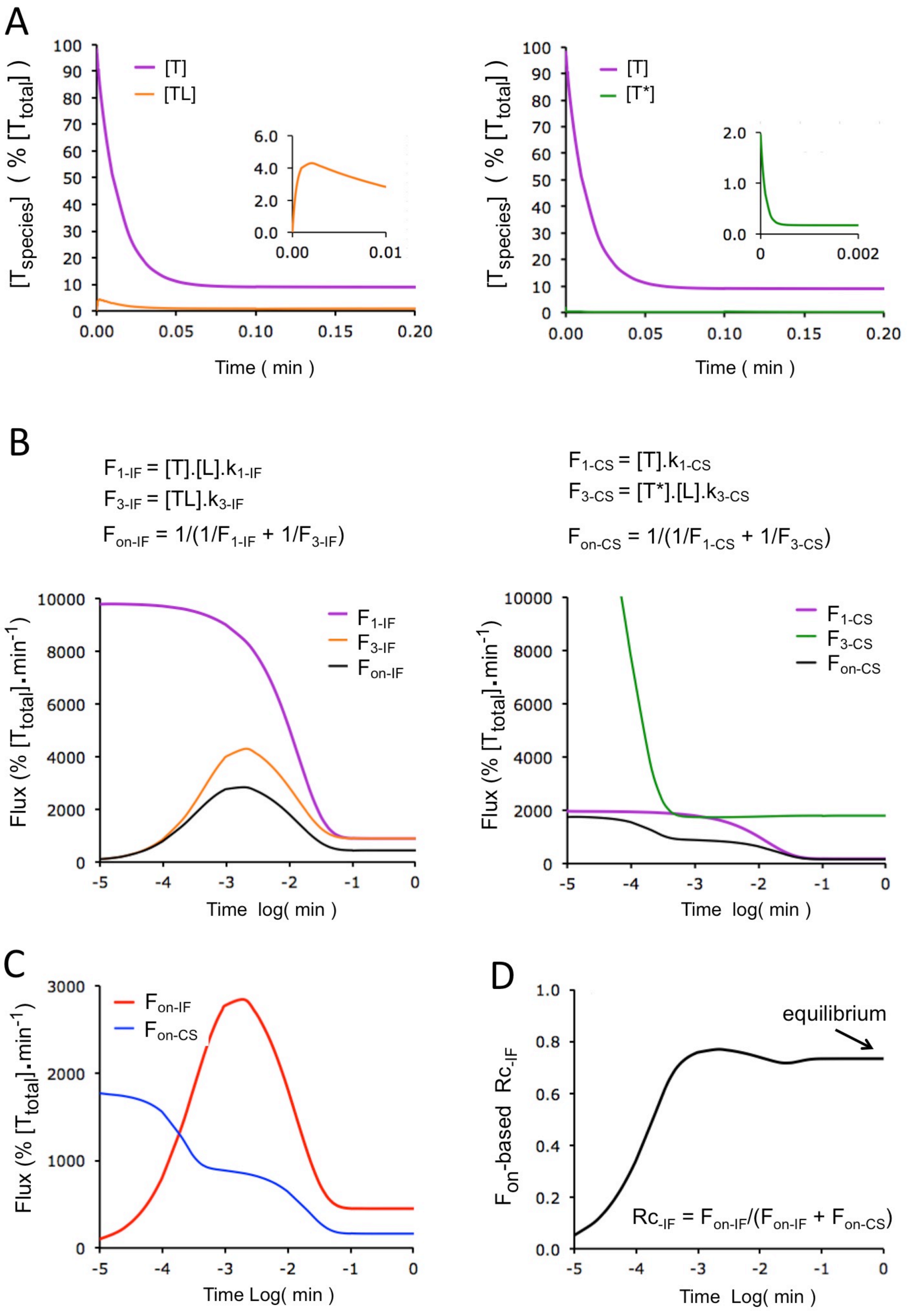
A
[T]
[TL]
[T*]
[Tspecies] ( % [Ttotal] )
Time ( min )
B
F1-IF = [T].[L].k1-IF
F3-IF = [TL].k3-IF
Fon-IF = 1/(1/F1-IF + 1/F3-IF)
F1-CS = [T].k1-CS
F3-CS = [T*].[L].k3-CS
Fon-CS = 1/(1/F1-CS + 1/F3-CS)
F1-IF
F3-IF
Fon-IF
F1-CS
F3-CS
Fon-CS
Flux (% [Ttotal]•min-1)
Time log( min )
C
Fon-IF
Fon-CS
Time Log( min )
D
equilibrium
Fon-based Rc-IF
Rc-IF = Fon-IF/(Fon-IF + Fon-CS)
Time Log( min )

**Figure S4**

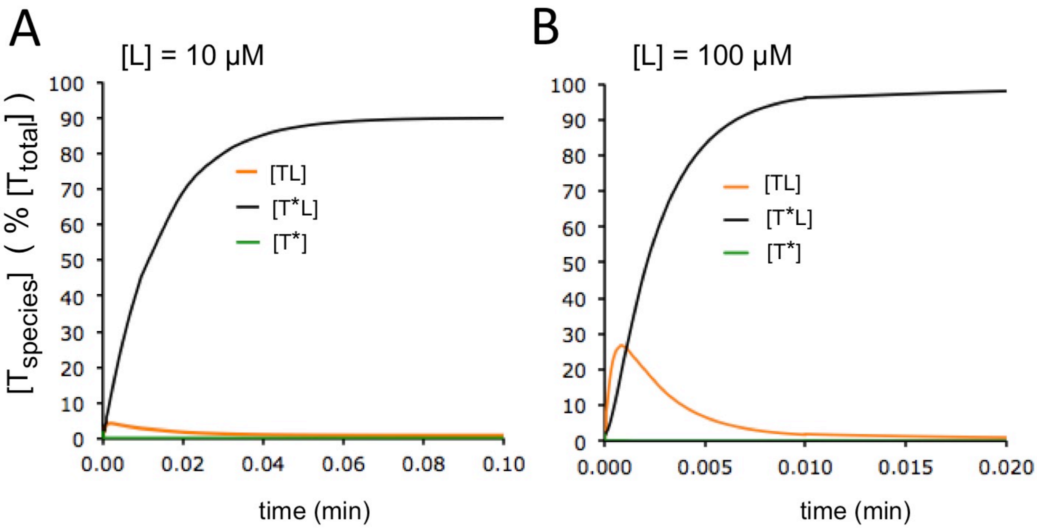


**Figure S5**

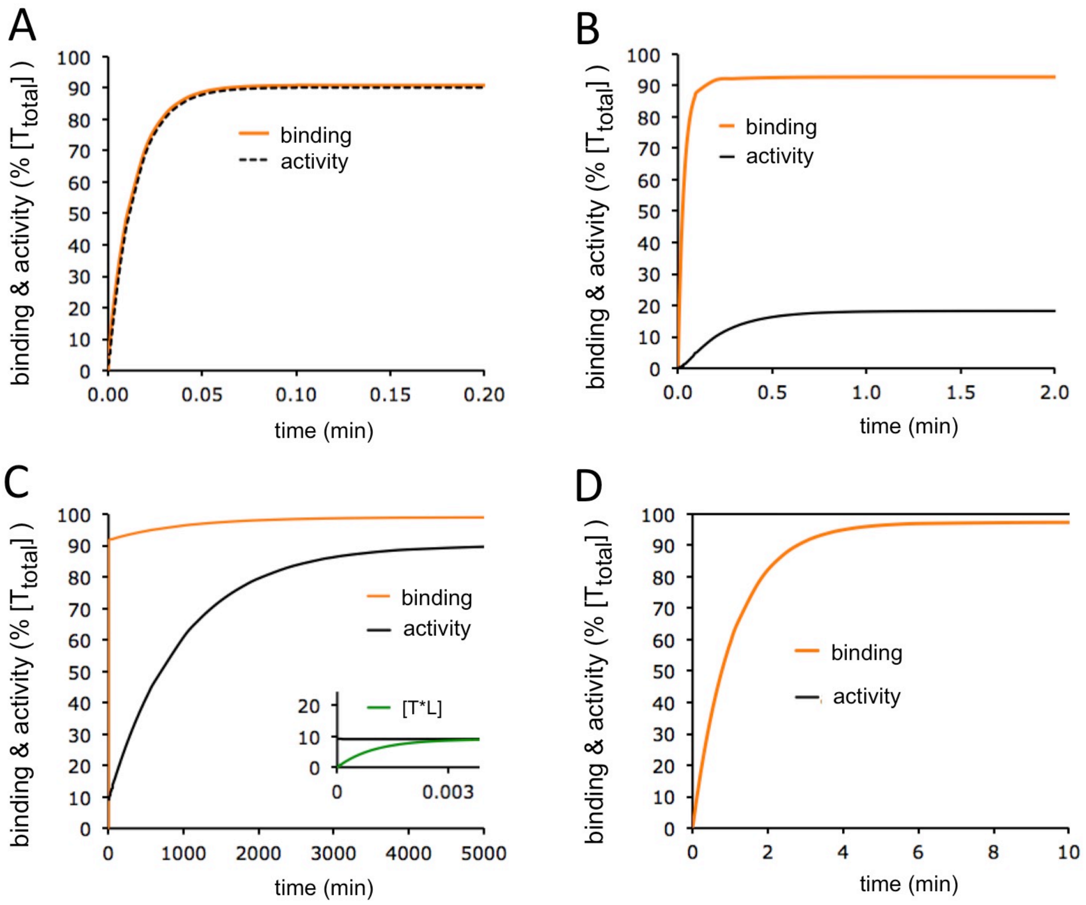

### Figure S6

Case A

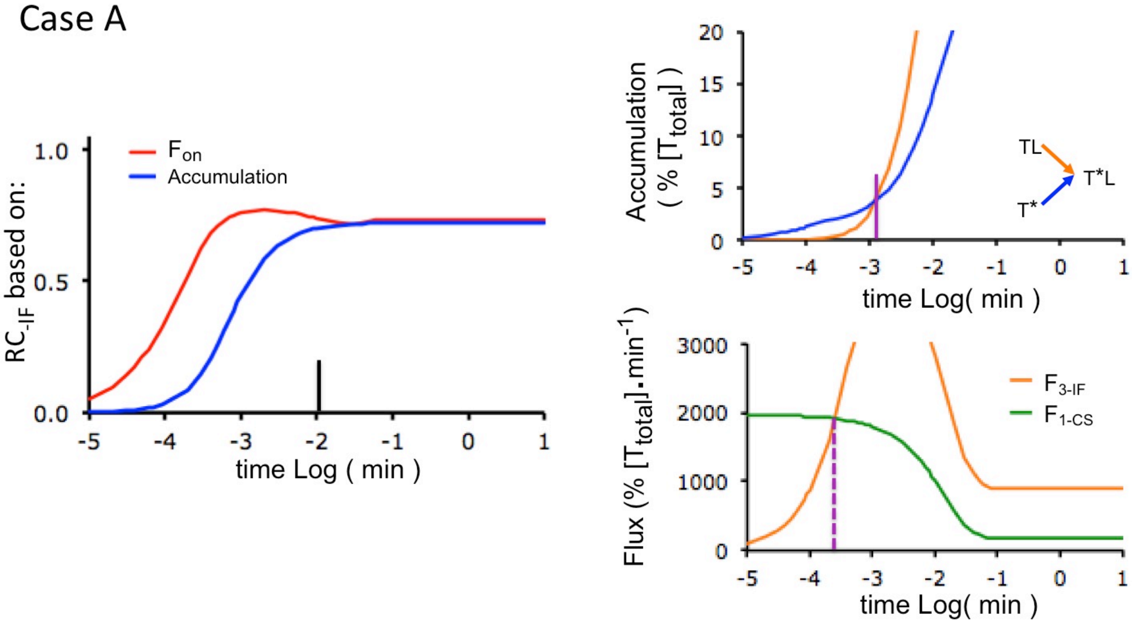


Case B

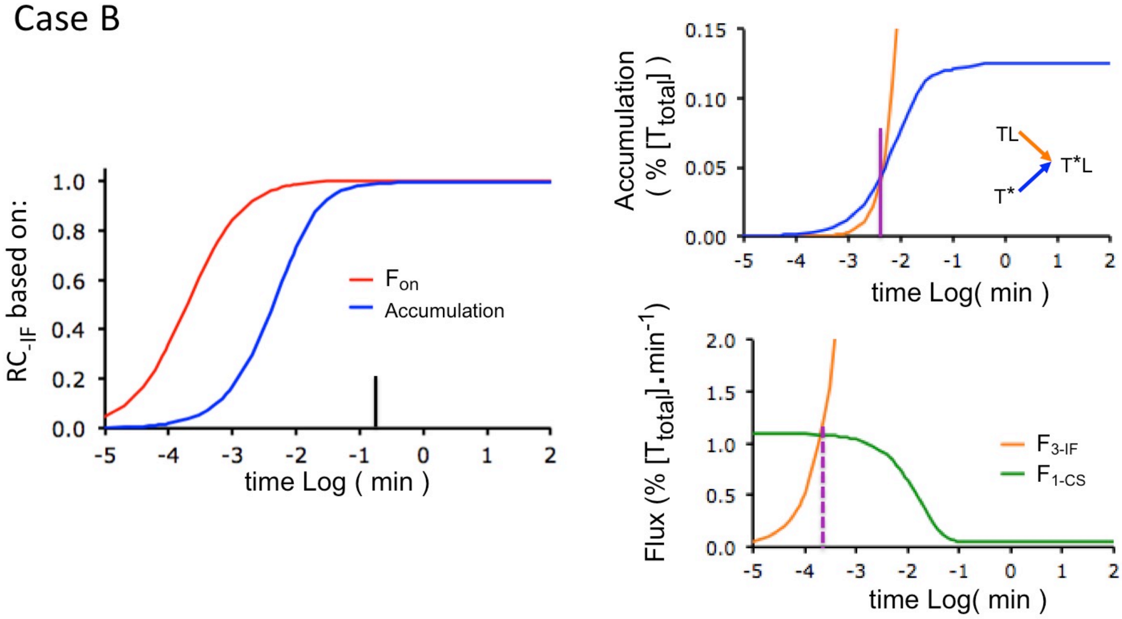

**Figure S7**

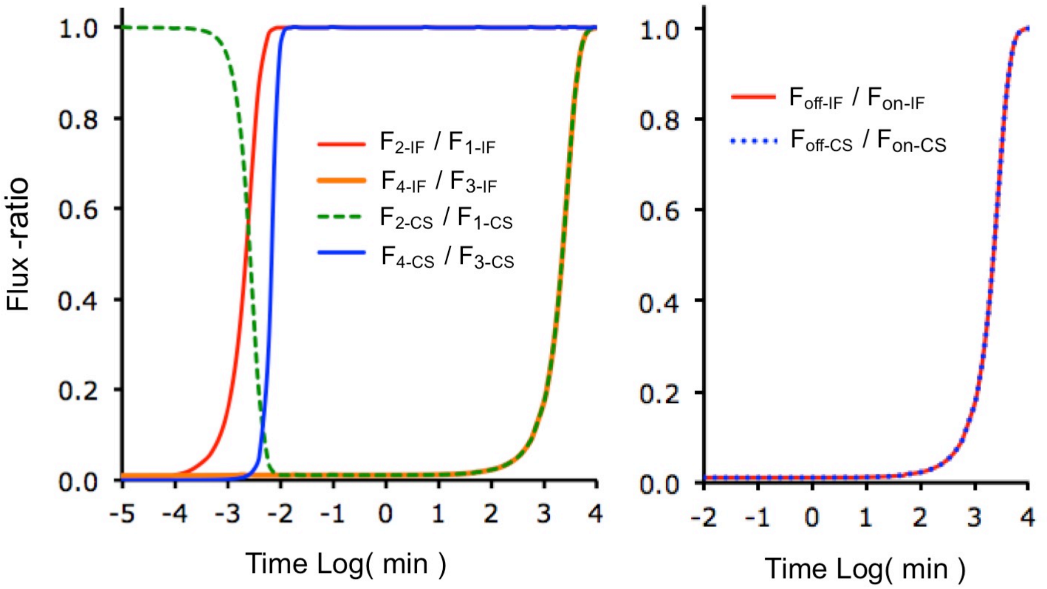


**Figure S8**

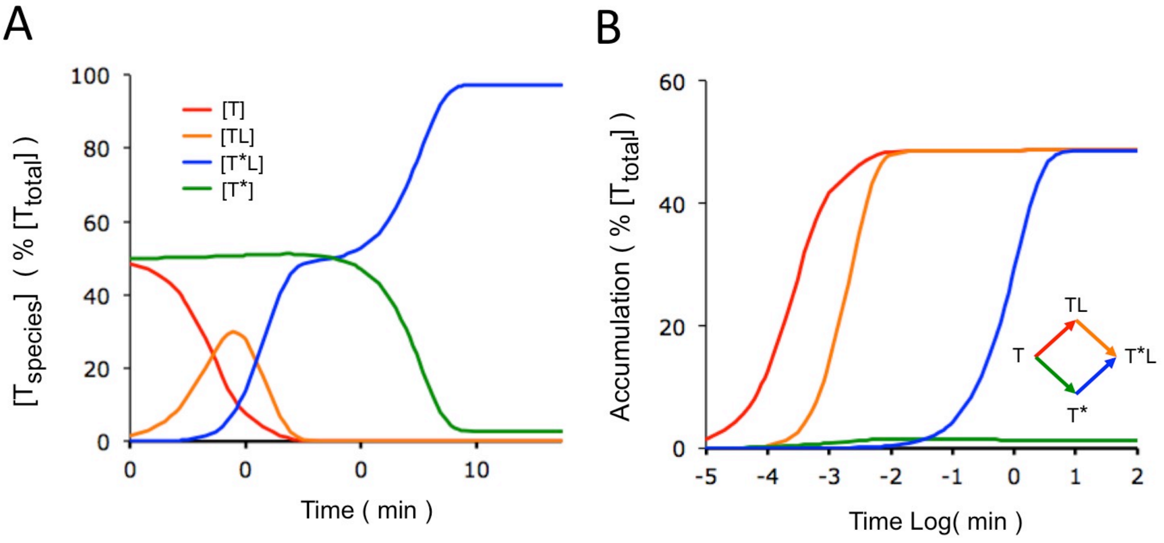

**Figure S9**

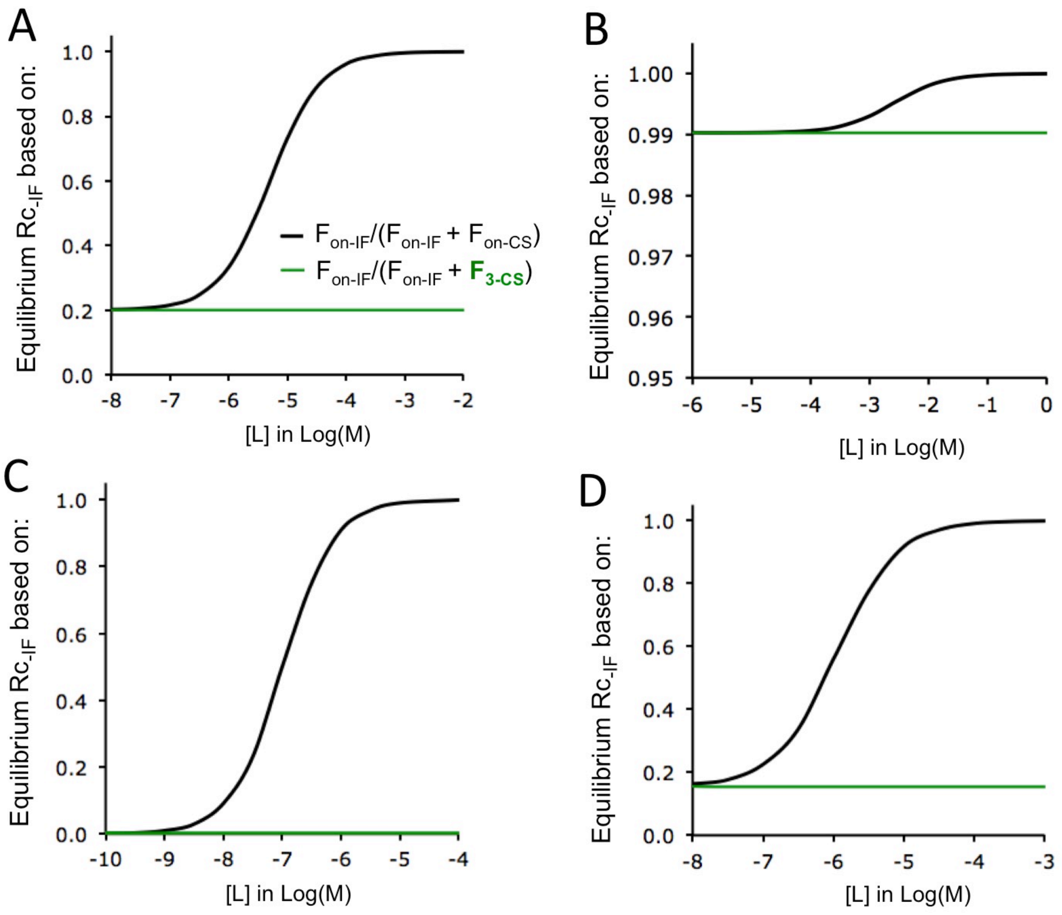
A
B
C
D
Equilibrium Rc-IF based on:
Fon-IF/(Fon-IF + Fon-CS)
Fon-IF/(Fon-IF + F3-CS)
[L] in Log(M)
1.0
0.8
0.6
0.4
0.2
0.0
-8
-7
-6
-5
-4
-3
-2
1.00
0.99
0.98
0.97
0.96
0.95
-1
0
-10
-9

#### Figure S10

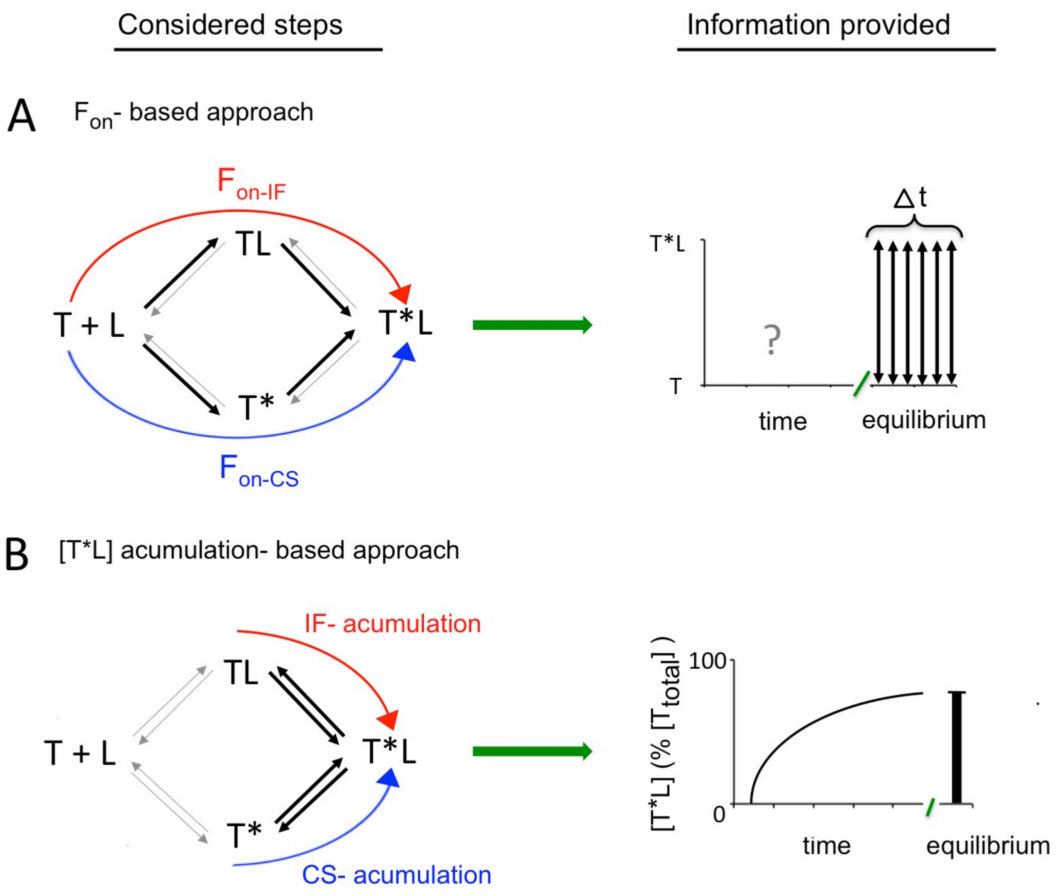
Considered steps
Information provided
A
Fon- based approach
Fon-IF
TL
T + L
T*L
T*
Fon-CS
Δt
T*L
?
T
time
equilibrium
B
[T*L] acumulation- based approach
IF- acumulation
TL
T + L
T*L
T*
CS- acumulation
[T*L] (% [Ttotal] )
100
0
time
equilibrium

**References:**


1) Hammes GG, Chang YC, Oas TG. Conformational selection or induced fit: a flux description of reaction mechanism. Proc Natl Acad Sci USA. 2009; 106:13737–13741. doi: 10.1073/pnas.0907195106. PMID: 19666553.

2) Zhou G, Pantelopoulos GA, Mukherrjee S, Voelz VA. Bridging microscopic and macroscopic mechanisms of p53-MDM2 binding with kinetic network models. Biophys J. 2017; 113: 785-793. doi: 10.1016/j.bpj.2017.07.009. PMID: 28834715.

3) Galburt EA, Rammohan J. Kinetic signature for parallel pathways: conformational selection and induced fit. Links and disconnects between observed relaxation rates and fractional equilibrium flux under pseudo-first-order conditions. Biochemistry. 2016; 55:7014-7022. doi: 10.1021/acs.biochem.6b00914. PMID: 27992996.

4) Michel D. Conformational selection or induced fit? New insights from old principles. Biochimie. 2016; 128-129:48–54. doi: 10.1016/j.biochi.2016.06.012. PMID: 27344613.

5) Tummino PJ, Copeland RA. Residence time of receptor-ligand complexes and its effect on biological function. Biochemistry. 2008; 47:5481–5492. doi: 10.1021/bi8002023. PMID: 18412369.

6) Vauquelin G. Distinct in vivo target occupancy by bivalent- and induced-fit-like binding drugs. Br J Pharmacol. 2017; 173:1268-1285 doi: 10.1111/bph.13989. PMID: 28838028

7) Vauquelin G, Maes D. Induced fit versus conformational selection: from rate constants to fluxes… and back to rate constants. Fund Res Perspect. 2021; 9: e00874. doi: 10.1002/prp2.847. PMID: 34459109.

8) Strickland S, Palmer G, Masset V. Determination of dissociation constants and specific rate constants of enzyme- substrate (or protein-ligand) interactions from rapid reaction kinetic data. J Biol Chem. 1975; 250:4048-4052. doi: https://doi.org/10.1016/S0021-9258(19)41384-7 PMID: 1126943.

9) Vogt AD, Di Cera E. Conformational selection or induced fit? A critical appraisal of the kinetic mechanism. Biochemistry. 2012; 51:5894-5902. doi: 10.1021/bi3006913. PMID: 22775458.

10) Vogt AD, Pozzi N, Chen Z, Di Cera E. Essential role of conformational selection in ligand binding. Biophys Chem. 2014; 186;13-21. doi: 10.1016/j.bpc.2013.09.003.
PMID: 24113284.

11) Lefkowitz RJ, Cotecchia S, Samama P, Costa T: Constitutive activity of receptors coupled to guanine nucleotide regulatory proteins. Trends Pharmacol Sci. 1993;14:303-307. doi: 10.1016/0165-6147(93)90048-O. PMID: 8249148.

12) Changeux JP, Edelstein S. Conformational selection or induced fit? 50 years of debate resolved. F1000 Biol Rep. 2011; 3:19. doi: 10.3410/B3-19. PMID: 21941598.

13) Kenakin T. New concepts in pharmacological efficacy at 7TM receptors: IUPHAR Review 2. Br J Pharmacol. 2012; 168:554-575. doi: 10.1111/j.1476-5381.2012.02223.x. PMID: 22994528.

14) Sekhar A, Velyvis A, Zoltsman G, Rosenzweig G, Bouvignies G, Kay LE. Conserved conformational selection mechanism of HP70 chaperone-substrate interactions. Elife. 2018; 7:e32764 doi: 10.7554/eLife.32764. PMID: 29460778.

15) Vauquelin G, Maes D. Competition in drug binding and … the race to equilibrium. Fund Clin Pharmacol. 2023; 37:147-157. doi : 10.1111/fcp.12824. PMID: 35981720.

16) Paton WDM. A theory of drug action based on rate of drug-receptor combination. Proc R Soc Lond B Biol Sci. 1961; 154:21-69. URL: https://www.jstor.org/stable/90247

17) Paton WDM. Kinetic theories of drug action with special reference to the acetyccholine group of agonists and antagonists. Ann NY Acad Sci. 1967;144:869-881. doi: 10.1111/j.1749-6632.1967.tb53816.x.

18) Clark A.J. General Pharmacology: in: 'Heffner's Handbuch Der Experimentellen

Pharmacologie Ergänzungsband, Band 4. Springer-Verlag: Berlin (1937).

19) Ariens EJ. Affinity and intrinsic activity in the theory of competitive inhibition. 1. Problems and theory. Arch Int Pharmacodyn Ther. 1954; 99:32-49. PMID: 13229418.

20) Stephenson RP. A modification of receptor theory Br J Pharmacol. 1956; 11:379-393. doi: 10.1111/j.1476-5381.1956.tb00006.x. PMID: 13383117.

21) Furchgott RF.. The use of β-haloalkylamines in the differentiation of receptors and in the determination of dissociation constants of receptor-agonist complexes. In: Harper NJ, Simmonds AB (eds). Advances in Drug Research., Vol. 3. Academic Press: New York, pp. (1966) 21-55.

22) Kenakin T. New concepts in pharmacological efficacy at 7TM receptors: IUPHAR Review 2. Br J Pharmacol. 2012; 168:554-575. doi: 10.1111/j.1476-5381.2012.02223.x. PMID: 22994528.

23) Leff P. The two-state model of receptor activation. Trends Pharmacol Sci. 1995; 16:89-97. doi: 10.1016/s0165-6147(00)88989-0. PMID: 754078.1

24) Colquhoun D. Binding, gating, affinity and efficacy: the interpretation of structure-activity relationships for agonists and of the effects of mutating receptors. Br J Pharmacol. 1998; 125:924-947. doi: 10.1038/sj.bjp.0702164. PMID: 9846630
